# Evaluating Coarse-Grained MARTINI Force-Fields for Capturing the Ripple Phase of Lipid Membranes

**DOI:** 10.1101/2020.12.02.408674

**Authors:** Pradyumn Sharma, Rajat Desikan, K. Ganapathy Ayappa

**Author notes:** Corresponding author: K. Ganapathy Ayappa. Co-first authors (Pradyumn Sharma, Rajat Desikan). These authors (P.S. and R.D.) contributed equally.

## Abstract

Phospholipids, which are an integral component of cell membranes, exhibit a rich variety of lamellar phases modulated by temperature and composition. Molecular dynamics (MD) simulations have greatly enhanced our understanding of phospholipid membranes by capturing experimentally observed phases and phase transitions at molecular resolution. However, the ripple (*P*_β′_) membrane phase, observed as an intermediate phase below the main gel-to-liquid crystalline transition with some lipids, has been challenging to capture with MD simulations, both at all-atom and coarse-grained (CG) resolution. Here, with an aggregate ~2.5 μs all-atom and ~122 μs CG MD simulations, we systematically assess the ability of six CG MARTINI 1,2-dipalmitoyl-sn-glycero-3-phosphocholine (DPPC) lipid and water force-field (FF) variants, parametrized to capture the DPPC gel and fluid phases, for their ability to capture the *P*_β′_ phase, and compared observations with those from an all-atom FF. Upon cooling from the fluid phase to below the phase transition temperature with smaller (380-lipid) and larger (> 2200-lipid) MARTINI and all-atom (CHARMM36 FF) DPPC lipid bilayers, we observed that smaller bilayers with both all-atom and MARTINI FFs sampled interdigitated *P*_β′_ and ripple-like states, respectively. However, while all-atom simulations of the larger DPPC membranes exhibited the formation of the *P*_β′_ phase, similar to previous studies, MARTINI membranes did not sample interdigitated ripple-like states at larger system sizes. We then demonstrated that the ripple-like states in smaller MARTINI membranes were kinetically-trapped structures caused by finite size effects rather than being representative of true *P*_β′_ phases. We showed that even a MARTINI FF variant that could capture the tilted *L*_β′_ gel phase, a prerequisite for stabilizing the *P*_β′_ phase, could not capture the rippled phase upon cooling. Our study reveals that the current MARTINI FFs (including MARTINI3) may require specific re-parametrization of the interaction potentials to stabilize lipid interdigitation, a characteristic of the ripple phase.

**TOC Graphic:** 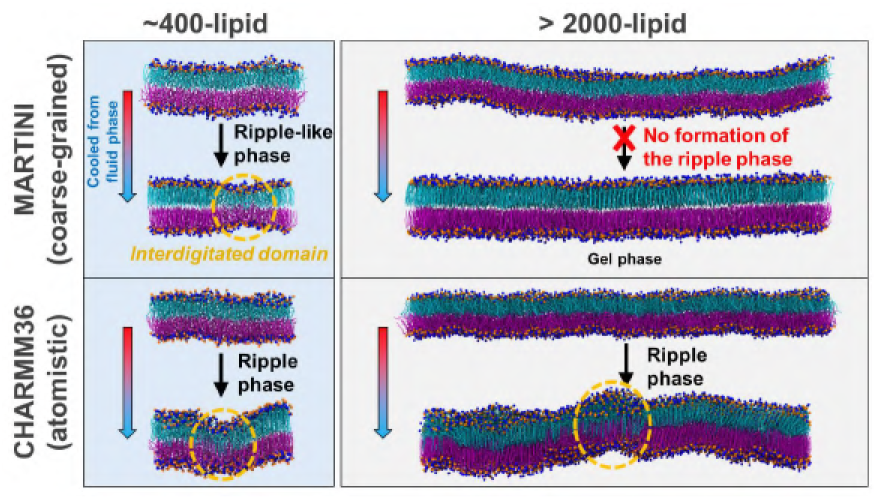

## Introduction

Lamellar lipid membranes, composed of two layers of amphiphilic lipids with solvated hydrophilic ‘heads’ and buried hydrophobic ‘tails’, exhibit a wide variety of thermotropic phases such as crystalline (*L*_c_), tilted gel (*L*_β′_) or straight gel (*L*_β_), tilted ripple (*P*_β′_), and liquid-crystalline (*L*_α_).^1,2^ Lipid bilayers are important constituents of biomembranes which are ubiquitous in living organisms^3^, and nanoparticle pharmaceuticals such liposomal drugs^4–6^. Specifically, lipid membrane phases influence: (i) biological processes at the cellular level such as membrane fusion and budding^7^, permeation of drugs^8,9^, effect of anaesthetics^10^, vulnerability to pathogenic toxins^11,12^, lipid flip-flop^13,14^, neural signalling^15^, and the function of membrane proteins^3,7^, and (ii) industrial applications such as the design, storage and usage of liposomal drugs^4–6,16^, cosmetic formulations^1^, and lipid-rich foods^1^. Molecular simulations of membranes at various levels of detail have yielded atomistic insights into the rich phase behaviour of phospholipid membranes while complementing experimental observations^1,17^. All-atom molecular dynamics (MD) simulations have successfully captured the low temperature *L*_β′_ or *L*_β_ phases, the liquid ordered and liquid disordered phases, and the main gel-to-fluid (*L*_β′_ to *L*_α_) phase transition in simple lipid bilayers while providing detailed insights into the rate-limiting formation of the melting seed that nucleates the transition^18^. Similarly, the molecular structure of lipid domains in the experimentally elusive *P*_β′_ phase, which commonly occurs upon cooling membranes from the *L*_α_ phase to below the gel-to-fluid transition temperature (pre-transition region), was unravelled at atomistic resolution via detailed MD simulations^19–21^ and was found to be consistent with experimental observations.

While these *in silico* studies illustrate the wide applicability of all-atom lipid force-fields (FFs), all-atom MD simulations are typically limited by computational bottlenecks to nanoscales, and complex membrane processes typically require microscale MD simulations^17,22^. Therefore, coarse-grained (CG) lipid models such as those used in the dissipative particle dynamics (DPD), MARTINI, and other frameworks have been developed^22–26^. Among the CG models, the MARTINI biomolecular FF has been successfully used to study a variety of complex biological systems outside the purview of all-atom MD simulations^22^. For example, MARTINI simulations have been used to carry out microsecond simulations of an idealized 20000-lipid plasma membrane comprising 63 different species of lipids^27^, a difficult feat with typical all-atom simulations. The MARTINI model is based on a four-to-one mapping scheme where four non-hydrogen atoms are typically modelled as effective CG interaction sites (‘beads’), whose parameters were calibrated by comparison with macroscopic properties such as thermodynamic transfer free energies (Details of bead types and FF parameterization in Refs.^22,24,28^). The versatility and computational tractability of CG models such as MARTINI render them as attractive options for capturing the microscopic details of complex membrane phenomena^22,24^. However, since CG models by definition possess reduced molecular detail^28^, their interaction potentials must be continuously assessed and refined, often by systematic comparison with experiments and/or corresponding detailed all-atom models^25,26,28–33^, to enable increasingly accurate CGMD simulations.

DPPC possesses a large headgroup and favourable inter-head interactions that stabilizes a tilted gel (*L*_β′_) phase at lower temperatures, unlike lipids with smaller headgroups such as phosphatidylethanolamine that favour the straight gel (*L*_β_) phase^20,21,34^. At higher temperatures, DPPC exhibits a main *L*_β′_ to *L*_α_ phase transition, with the *P*_β′_ phase occurring at pre-transition temperatures^34–40^. The *P*_β′_ phase has interesting properties such as the influence of thermal history on the symmetry of the ripple domains^20^, and therefore has been the subject of extensive experimental^35,39–42^ and theoretical^18–21,25,31,37,38,43–47^ studies. In Table 1, we summarize some of the previous studies where the rippled phase has been observed, including details of lipid type, resolution (all-atom, united atom or coarse-grained) and FF. Despite extensive *in silico* investigations, MD simulations that capture the *P*_β′_ phase have proven to be challenging since the ripple domain structure and formation is not only a function of the specific lipid FF (Table 1), but also depends on the cooling/heating protocols^18,36^ and system size effects^20,36,45^. In spite of these challenges, MD simulations of lipid bilayers in the ripple phase have revealed elusive details of molecular organization that were challenging to probe via experiments, such as the asymmetric sawtooth-like molecular organization of lipids in the DPPC ripple domains^19–21^.

**Table 1.** List of previous studies which have explored the *P*_β′_ phase using MD simulations.

| *Lipid* | *Year* | *Resolution* | *Force-field* | *Comparison with experiments* | *Citation* |
| --- | --- | --- | --- | --- | --- |
| DPPC | 2005 | United atom | Modified Berger lipid | Qualitative comparison with DMPC electron density profiles | Ref.<sup>19</sup> |
| DMPC | 2005 | Coarse-grained | DPD | – | Ref.<sup>21</sup> |
| DPPC | 2007 | Coarse-grained | Custom (single tailed) | Qualitative comparison with DPPC electron density maps | Ref.<sup>20</sup> |
| DMPC, DPPC | 2012 | Coarse-grained | DPD, MARTINI | Enthalpic signatures of phase transitions from MD compared with differential scanning calorimetry | Ref.<sup>36</sup> |
| BTMAC+SA | 2014 | United atom | GROMOS 87, OPLS | – | Ref.<sup>43</sup> |
| DPPC | 2014 | Coarse-grained | MARTINI | – | Ref.<sup>48</sup> |
| DMPC | 2016 | All-atom | CHARMM36, Lipid14 | – | Ref.<sup>44</sup> |
| DPPC | 2017 | Coarse-grained | Dry MARTINI | – | Ref.<sup>49</sup> |
| DMPC, DPPC | 2018 | All-atom | CHARMM36 | Comparison of bilayer properties with X-ray diffraction experiments | Ref.<sup>46</sup> |
| DPPC | 2018 | All-atom | CHARMM36 | – | Ref.<sup>18</sup> |
| DPPC | 2018 | All-atom, coarse-grained | CHARMM36, MARTINI | – | Ref.<sup>31</sup> |
| DPPC | 2019 | All-atom | CHARMM36 | – | Ref.<sup>45</sup> |
| BTMAC+SA | 2020 | All-atom, coarse-grained | CHARMM36, MARTINI | – | Ref.<sup>25</sup> |

Abbreviations: DPD – dissipative particle dynamics; DPPC – 1,2-dipalmitoyl-sn-glycero-3-phosphocholine; DMPC – 1,2-dimyristoyl-sn-glycero-3-phosphocholine; BTMAC – benzyltrimethylammonium chloride; SA – stearyl alcohol.

While DPPC membranes described by the standard MARTINI FF and its variants have been shown to capture the gel and fluid states^22,31^, there are conflicting reports of their ability to capture the ripple phase. Some MARTINI studies have reported ripple formation both with and without enhanced sampling^31,48,49^, while others have reported that MARTINI membranes cannot capture the ripple phase^36^, with the interdigitated ripple-like domains observed upon quasi-static cooling from the *L*_α_ phase reported to be artefacts caused by finite size effects^36^. Therefore, the ability of MARTINI membranes to capture the *P*_β′_ phase still remains an unresolved problem. To systematically address this issue, we assess the ability of commonly employed^18,19,23,29,31,36,45,46,48^ DPPC bilayers, described by six MARTINI lipid FF variants (described subsequently), to capture the interdigitated tilted ripple membrane phase (*P*_β′_).

We first establish a standard simulated-annealing-based cooling MD protocol, built on previous studies^18–20,31,45,46^, that consistently yielded a ripple phase upon cooling from the fluid phase to below the gel-to-fluid phase transition temperature with all-atom DPPC membranes across multiple simulation replicates. By employing the same protocol with MARTINI DPPC membranes, and by fixing other important variables such as initial configurations (*L*_α_ phase), system sizes and cooling rates, we checked whether the MARTINI membranes yielded stable ripple phase structures, similar to those observed in all-atom simulations. Six MARTINI lipid and water FF combinations were tested: (1) standard MARTINI version 2.2 lipid FF with normal water (‘MARTINI2n’)^23^, or polarizable water^50^ (‘MARTINI2p’) with embedded charges and explicit charge screening for better electrostatics^22^, (2) optimized MARTINI FF for DPPC – has refined bond parameters for better reproduction of DPPC membrane phase behaviour and can capture the tilted *L*_β′_ gel phase – with either normal (‘MARTINI2On’) or polarizable (‘MARTINI2Op’) water, and, (3) the recently released MARTINI 3 alpha (‘MARTINI3a’)^33^, and the slightly different previous interim version MARTINI 3 beta (version 3.0.b.3.2; ‘MARTINI3b’)^51^, with their in-built water models. The next generation MARTINI3 FFs include new parameterization of charged beads as well as small and tiny beads involved in aromatic groups, improved bead interactions, and extensive benchmarking, and MARTINI3a is poised to replace MARTINI2 as the default choice of MARTINI FF^33,51^. For each FF variant, 10 independent replicates of simulated cooling were performed, ensuring sufficient sampling for robust statistical estimates^18,52^. We observed that smaller (380-lipid) MARTINI DPPC bilayers sampled interdigitated ripple-like states that were not observed with larger DPPC membranes (> 2200-lipid). The ripple-like states in these smaller MARTINI DPPC membranes were demonstrated to be kinetically-trapped structures caused due to finite size effects, rather than being representative of a true ripple phase. However, all-atom simulations of both the smaller and larger DPPC membranes exhibited the formation of the *P*_β′_ phase, similar to previous studies. Our observations thus assess the ability of MARTINI lipid FFs to capture the ripple phase and serve as a guide for future CG lipid FF development.

## Methods

### All-atom MD simulations of DPPC membranes

The MD protocols employed in this study for simulating DPPC membranes in the gel, ripple and fluid phases are briefly described below. All the simulations performed in this study are summarized in Table 2.

**Table 2.** Summary of all-atom and MARTINI simulations performed in this study.

| Forcefield | Time (μs)<sup>#</sup> | Number of replicas | System size (L<sub>x</sub> × L<sub>y</sub> × L<sub>z</sub> nm<sup>3</sup>) | Number of lipids | Number of particles | Type of MD simulation<sup>a</sup> | Temp. (K) | Figures |
| --- | --- | --- | --- | --- | --- | --- | --- | --- |
| *Small membrane simulations* |  |  |  |  |  |  |  |  |
| CHARMM36 | 0.10 | 1 | 15.7 × 7.8 × 8.4 | 368 | 103040 | EQMD<sup>a</sup> | 350 | 1, S1 |
| CHARMM36 | 0.36 | 2 | 13.4 × 6.7 × 10.6 | 368 | 103040 | SAMD | 350-290 |  |
| CHARMM36 | 0.20 | 2 | 13.4 × 6.7 × 10.6 | 368 | 103040 | EQMD | 290 |  |
| MARTINI2n | 0.10 | 1 | 15.8 × 7.9 × 9.6 | 380 | 9626 | EQMD<sup>a</sup> | 350 | 2 |
| MARTINI2On | 0.10 | 1 | 15.8 × 7.9 × 9.6 | 380 | 9620 | EQMD<sup>a</sup> | 350 | - |
| MARTINI3a | 0.10 | 1 | 16.1 × 8.0 × 8.8 | 380 | 9626 | EQMD<sup>a</sup> | 350 | - |
| MARTINI3b | 0.10 | 1 | 15.9 × 7.9 × 9.1 | 380 | 9626 | EQMD<sup>a</sup> | 350 | - |
| MARTINI2p | 0.10 | 1 | 16.0 × 8.0 × 9.0 | 380 | 19761 | EQMD<sup>a</sup> | 350 | - |
| MARTINI2Op | 0.10 | 1 | 15.5 × 7.8 × 9.5 | 380 | 19764 | EQMD<sup>a</sup> | 350 | - |
| MARTINI2n | 0.42 | 10 | 14.2 × 7.1 × 11.1 | 380 | 9626 | SAMD | 350-280 | S2 |
| MARTINI2On | 0.42 | 10 | 13.9 × 7.0 × 11.5 | 380 | 9620 | SAMD | 350-280 | S3 |
| MARTINI3a | 0.42 | 10 | 14.1 × 7.1 × 10.5 | 380 | 9626 | SAMD | 350-280 | S4 |
| MARTINI3b | 0.42 | 10 | 14.2 × 7.1 × 10.5 | 380 | 9626 | SAMD | 350-280 | S5 |
| MARTINI2p | 0.42 | 10 | 14.4 × 7.2 × 10.2 | 380 | 19761 | SAMD | 350-280 | S6 |
| MARTINI2Op | 0.42 | 10 | 13.8 × 6.9 × 10.9 | 380 | 19764 | SAMD | 350-280 | S7 |
| MARTINI2n | 5 | 2<sup>b</sup> | 14.3 × 7.1 × 11.0 | 380 | 9626 | EQMD | 280 | 2, 6 |
| MARTINI2On | 5 | 3<sup>c</sup> | 13.9 × 7.0 × 11.4 | 380 | 9620 | EQMD | 280 | 2, 6, S13 |
| MARTINI3a | 5 | 1<sup>d</sup> | 14.1 × 7.1 × 10.5 | 380 | 9626 | EQMD | 280 | 2 |
| MARTINI3b | 5 | 1<sup>e</sup> | 14.0 × 7.0 × 10.7 | 380 | 9626 | EQMD | 280 | S5b |
| MARTINI2p | 5 | 1<sup>f</sup> | 14.3 × 7.2 × 10.2 | 380 | 19761 | EQMD | 280 | 2 |
| MARTINI2Op | 5 | 1<sup>g</sup> | 13.8 × 6.9 × 10.9 | 380 | 19764 | EQMD | 280 | 2 |
| *Large membrane simulations* |  |  |  |  |  |  |  |  |
| CHARMM36 | 0.02 | 1 | 46.4 × 15.4 × 16.1 | 2208 | 1135872 | EQMD<sup>a</sup> | 350 | 3, S8 |
| CHARMM36 | 0.36 | 2 | 40.3 × 13.4 × 19.9 | 2208 | 1135872 | SAMD | 350-290 |  |
| CHARMM36 | 0.20 | 2 | 40.4 × 13.5 × 19.8 | 2208 | 1135872 | EQMD | 290 |  |
| MARTINI2n | 0.10 | 2 | 46.5 × 15.5 × 20.6 | 2204 | 112719 | EQMD<sup>a</sup> | 350 | 4 |
| MARTINI2On | 0.10 | 2 | 46.3 × 15.4 × 20.8 | 2204 | 112719 | EQMD<sup>a</sup> | 350 | - |
| MARTINI3a | 0.10 | 2 | 46.6 × 15.5 × 19.1 | 2204 | 112717 | EQMD<sup>a</sup> | 350 | - |
| MARTINI3b | 0.10 | 2 | 47.1 × 15.7 × 18.8 | 2204 | 112717 | EQMD<sup>a</sup> | 350 | - |
| MARTINI2n | 0.42 | 2 | 39.9 × 13.3 × 26.0 | 2204 | 112719 | SAMD | 350-280 | 4, S10 |
| MARTINI2On | 0.42 | 2 | 39.9 × 13.1 × 26.6 | 2204 | 112719 | SAMD | 350-280 |  |
| MARTINI3a | 0.42 | 2 | 39.3 × 13.1 × 24.9 | 2204 | 112717 | SAMD | 350-280 |  |
| MARTINI3b | 0.42 | 2 | 41.3 × 13.8 × 22.7 | 2204 | 112717 | SAMD | 350-280 |  |
| MARTINI2n | 3 | 1 | 39.6 × 13.2 × 12.7 | 2280 | 57756 | EQMD | 280 | 5, S12 |
| MARTINI2On | 5, 8, 10 | 3 | 39.7 × 13.2 × 12.6 | 2280 | 57720 | EQMD | 280 | 5, 6, S12, S13 |
| MARTINI3a | 10 | 1 | 39.8 × 13.3 × 11.9 | 2280 | 57756 | EQMD | 280 | 5, S12 |
| MARTINI3b | 10 | 1 | 40.0 × 13.3 × 11.8 | 2280 | 57756 | EQMD | 280 | S12 |
| MARTINI2On | 0.10 | 1<sup>h</sup> | 46.5 × 46.5 × 20.2 | 6728 | 333604 | EQMD<sup>a</sup> | 350 | S11 |
| MARTINI2On | 0.42 | 2<sup>h</sup> | 39.5 × 39.5 × 26.0 | 6728 | 333604 | SAMD | 350-280 | S11 |

<sup>#</sup>Simulation times from MARTINI CGMD were not scaled. Total simulation time in this study: ~2.5 μs (all-atom MD) and ~122 μs (MARTINI CGMD). <sup>a</sup>Initial simulations to equilibrate starting configurations. <sup>a</sup>EQMD – equilibrium MD simulations in the NPT ensemble; SAMD – simulated annealing MD simulations at constant pressure. <sup>b</sup>Replicas with the ‘best’ ripple-like (Repl. 1 in Figure S2) and straight gel (Repl. 7 in Figure S2) phases were further equilibrated for 5000 ns. <sup>c</sup>Replicas with the ‘best’ ripple-like (Repl. 5 in Figure S3), tilted ripple-like (Repl. 7 in Figure S3) and tilted gel (Repl. 3 in Figure S3) phases were further equilibrated for 5000 ns. Replicas with the ‘best’ ripple-like phase: <sup>d</sup>Repl. 5 in Figure S4, <sup>e</sup>Repl. 9 in Figure S5, <sup>f</sup>Repl. 5 in Figure S6, and <sup>g</sup>Repl. 10 in Figure S7, were further equilibrated for 5000 ns each. <sup>h</sup>Square membrane patch; all others are rectangular membrane patches.

For the small membrane atomistic simulations, two charge-neutral, solvated, fluid phase (*L*_α_), 368-lipid membrane configurations at 350 K, with similar spatial coordinates but different starting velocities, were set up using CHARMM-GUI^53–55^ as independent replicates (Figures 1 and S1). The size of the lipid patches was approximately 15.2 nm (x-axis) × 7.6 nm (y-axis) × 9.4 nm (z-axis), with the characteristic dimension of the simulation box in the membrane plane in the range of the periodicity of 12-16 nm corresponding to previously observed ripple profiles in DPPC membranes^19,56^. This was important because the quality of the observed ripples was found to depend on the characteristic box dimension along the ripple formation^20^. Systems whose characteristic box length in the membrane plane were non-integer multiples of ripple periodicity resulted in frustrated and interconnected structures, unlike the clean ripples observed in systems with integer multiple box lengths^20^. Each system contained >10^5^ atoms. The water:lipid ratio was set at 50:1, ensuring sufficient hydration normal to the membrane plane (z), so that the systems could be approximated as isolated lipid bilayers along the z-axis.

**Figure 1.**
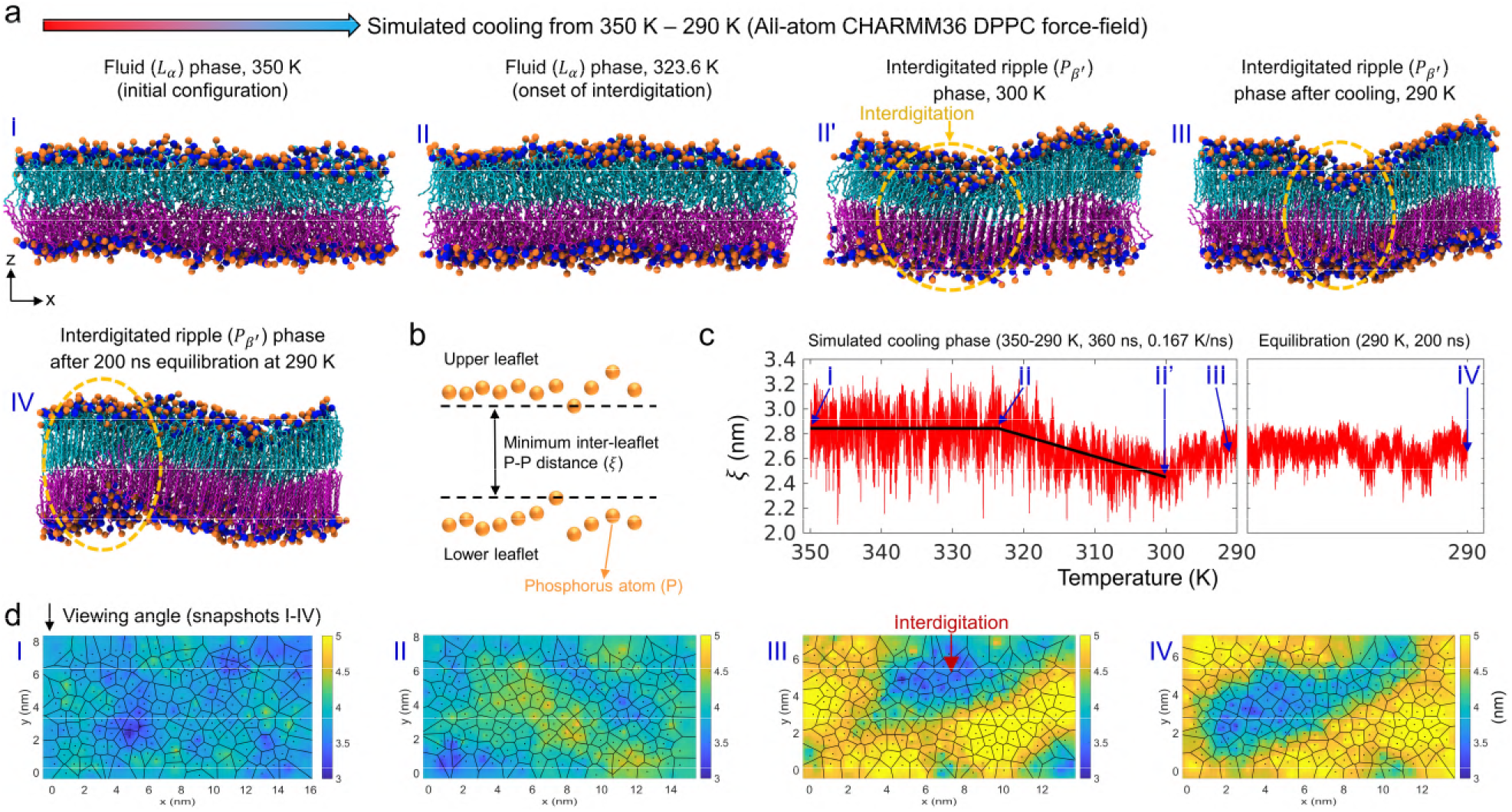
All-atom Lα DPPC membranes consistently form ripples upon cooling. **(a)** MD snapshots (colouring described in Methods) during the cooling of a 368-lipid, all-atom, DPPC membrane from 350 K to 290 K. Time points: I – initial equilibrated *L*_α_ phase membrane at 350 K before the start of cooling, II – start of interdigitation, IIʹ –time at which the average inter-leaflet distance in the time series is at a minimum (300 K), III – end of simulated cooling (290 K), and IV – end of equilibration MD at constant temperature (290 K). Interdigitated domains are highlighted by yellow dashed circles. **(b)** Schematic illustrating the minimum inter-leaflet P-P distance, ξ, which is used as an order parameter to track leaflet interdigitation (Equation 1). **(c)** Time trace (red) and biphasic linear fits (black) of ξ in cooling simulations of atomistic DPPC membranes. **(d)** Voronoi diagram from the x-y coordinates of the upper leaflet P atoms depicting the area per head group of individual lipids is superimposed on the thickness modulations of membranes at time points I-IV, corresponding to the MD snapshots in (a) (viewing angle pictorially depicted).

Atomistic simulations were performed using the CHARMM36 FF^44,57,58^ implementation (March 2019 version from http://mackerell.umaryland.edu/charmm_ff.shtml) in Gromacs (version 2018.6)^59^, according to the currently advocated best practices for lipid bilayer simulations^17,60–62^. (See Refs.^30,44^ for a brief discussion of all-atom and united atom lipid FFs.) A leapfrog integrator (2 fs integration time step) was used along with Verlet buffered lists and three-dimensional periodic boundary conditions. Electrostatic interactions were computed with the particle mesh Ewald algorithm^63^, and van der Waal interactions were smoothly switched to zero between 1.0 to 1.2 nm. Covalent bonds involving hydrogen atoms were constrained using the LINCS algorithm^64^. Simulations were performed in the NPT ensemble with semi-isotropic pressure coupling along the lateral bilayer plane, by employing the stochastic velocity-rescaling thermostat^65^ for temperature control, and the Parrinello-Rahman barostat^66^ for pressure control, with coupling constants of 0.1 and 2.0 ps, respectively. The isothermal compressibilities for the barostat were *k_XY_* = *k_Z_* = 4.5 × 10^−5^ bar^−1^. After energy minimization and equilibration (100 ns) of the initial configurations in the *L*_α_ phase (350 K) according to standard CHARMM-GUI protocols^54^, the systemic temperatures of the replicates were continuously reduced from 350 K to 290 K at the rate of 1 K per 6 ns, or −0.167 K/ns, by employing the stochastic simulated annealing MD (SAMD) algorithm implemented in Gromacs^59^ (360 ns per SAMD replicate). We chose to cool slightly beyond the pre-transition temperature of 300-310 K at which the *P*_β′_ phase is typically observed^18,34,36–38^, because super-cooling is often required in kinetic MD simulations for eliciting a phase transition^18,67,68^. The resulting cooled bilayers at 290 K were further equilibrated at a constant temperature of 290 K for 200 ns for thermal and structural stability.

To test for finite size effects and to compare with MARTINI simulations (described next), two large 2208-lipid membranes measuring 47.0 nm × 15.7 nm × 15.0 nm and possessing a high water:lipid ratio of 128:1 (>10^6^ atoms total; Table 2) were equilibrated in the *L*_α_ phase at 350 K. Subsequently, both replicates were cooled to 290 K at the rate of −0.167 K/ns (chosen based on Ref.^18^) and then equilibrated at 290 K for an additional 200 ns.

### MARTINI simulations of DPPC membranes

380-lipid MARTINI DPPC membranes of size 15 nm × 7.5 nm × 10 nm were prepared using the ‘INSANE’ python module^69^. The water-bead:lipid ratio was ~13.3:1. Since each water bead is equivalent to 4 water molecules, this corresponds to a water:lipid ratio of ~53:1, similar to the small all-atom membranes. As described in the Introduction, six MARTINI FF variants were used in this study: MARTINI2n^23^, MARTINI2p^50^, MARTINI2On^31^, MARTINI2Op^31,50^, MARTINI3a^33^, and MARTINI3b^51^. MARTINI2n and MARTINI2On systems also contained anti-freeze solvent beads^23^ to prevent any artefactual freezing of MARTINI water beads.

Input CGMD parameters were fixed based on previously reported parameter settings for improved performance^70^. All CGMD simulations were performed with a 20 fs timestep in the NPT ensemble by employing the stochastic velocity-rescaling thermostat^65^ (coupling constant of 1.0 ps) and the Parrinello-Rahman semi-isotropic barostat^66^ (coupling constant of 12.0 ps). The isothermal compressibilities for the barostat were *k_XY_* = *k_Z_* = 3 × 10^−4^ bar^−1^. Electrostatics were treated with the reaction-field algorithm^71^ with a cut-off of 1.1 nm, relative dielectric constant of 2.5 and 15 for systems with and without polarizable water, respectively, and the relative dielectric constant of the reaction field set to infinity. The van der Waal forces were computed with a cut-off of 1.1 nm. Post energy minimization, six CGMD membranes, each described by a MARTINI FF variant, were equilibrated in the *L*_α_ phase at 350 K for 100 ns, followed by simulated cooling from 350 K to 280 K at −0.167 K/ns (420 ns per SAMD replicate). For each FF, 10 independent SAMD replicates were simulated for ensuring robustness of observations^52^, resulting in a total of 60 CG-SAMD simulations (Figures 2, S2-S7). The replicates which yielded the best ripple-like phases were further equilibrated for 1-5 μs at 280 K to ensure that the ripples had attained thermal and structural equilibrium (Figures 2 and S5*b*).

**Figure 2.**
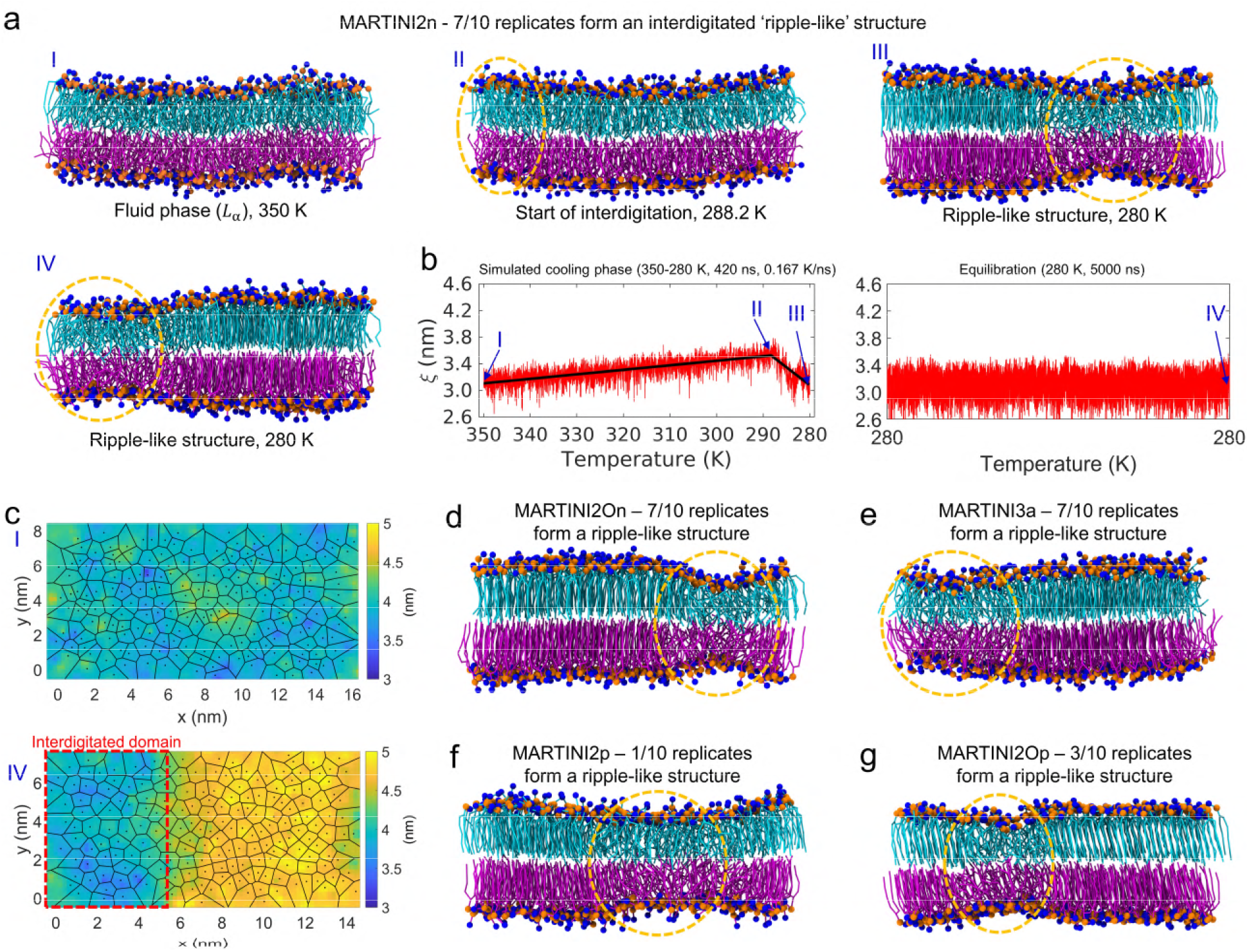
Smaller MARTINI membranes form a ripple-like interdigitated phase upon cooling. **(a)** Similar to all-atom MD (Figure 1), snapshots from time points I-IV from the simulated cooling of a MARTINI2n membrane replicate from 350 K – 280 K are shown (Replicate 1 from Figure S2). Interdigitated regions are highlighted by a yellow dashed circle. **(b)** Minimum inter-leaflet P-P distance with time in the 420 ns cooling and 5000 ns constant temperature equilibration phases. **(c)** Voronoi-thickness maps contrasting the initial fluid membrane and the final interdigitated ripple-like membrane (time points I and IV, respectively). Cooling of 7/10 independent MARTINI2n MD replicates resulted in the formation of a ripple-like phase (Figure S2). Other MARTINI FF variants: **(d)** MARTINI2On (Replicate 5 from Figure S3), **(e)** MARTINI3a (Replicate 5 from Figure S4; MARTINI3b in Figure S5), **(f)** MARTINI2p (Replicate 5 from Figure S6; formed a ripple-like state during EQMD and not SAMD) and **(g)** MARTINI2Op (Replicate 10 from Figure S7), show the formation of ripple-like phases in 7, 7, 1 and 3 out of 10 replicates each, respectively. These snapshots correspond to time point IV.

To test for finite-size effects, many large MARTINI DPPC bilayer configurations, with both rectangular and square geometries, were constructed and simulated (Table 2).

### Analysis and visualization

MD snapshots are illustrated using the visual molecular dynamics 1.9.3 (VMD)^72^ software. The DPPC tails in the upper and lower leaflets of both all-atom and MARTINI membranes are consistently colored cyan and magenta, respectively. In all-atom membranes, phosphorus atoms (P) are colored orange, and nitrogen atoms in the choline group are colored blue. Similarly, in MARTINI membranes, the phosphate bead is colored orange and the choline bead is colored blue. The minimum inter-leaflet P-P distance (ξ), which is an order parameter to track leaflet interdigitation (Figure 1*b*), was computed using the inbuilt ‘gmx mindist’ tool in Gromacs. The ξ from the fluid and interdigitated regimes are fit to a four-parameter piecewise-linear function (Equation 1) to accurately estimate the time of the onset of membrane interdigitation, τ.

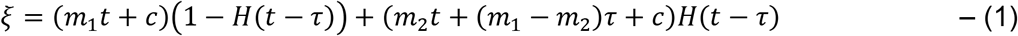

Here, *m*_1_ and *m*_2_ are the slopes of the two lines corresponding to the fluid and interdigitated regimes (when it occurs), respectively, *c* is the intercept of the fluid-phase line (the intercept of the other line is (*m*_1_ − *m*_2_)τ + *c* from the point of intersection), and *H*(*t* − τ) is the Heaviside function which is zero for *t* < τ, and one for *t* ≥ τ. If interdigitation occurs, *m*_2_ < 0, else *m*_2_ > 0 (illustrated in Figure S2). The fitting was performed using the ‘lsqcurvefit’ function (optimization toolbox) in conjunction with ‘multistart’ (global optimization toolbox) implemented in MATLAB 2018a (https://www.mathworks.com). All other plots are also created with MATLAB. The combined Voronoi-thickness modulation heatmaps are plotted using the ‘voronoi’ MATLAB function on the lipid phosphorus atom x, y coordinates (upper leaflet) from the corresponding MD snapshot. The membrane thickness^43^ is computed using the ‘Membplugin’ 1.1 tool^73^ in VMD and visualized using MATLAB. Similar to a previous study^74^, lipid tilt angles from MARTINI membranes (*L*_β_ vs. *L*_β′_) were computed from the last 10 ns of the respective MD trajectories, using the inbuilt ‘gmx gangle’ tool in Gromacs. The lipid tilt was defined as the angle between the lipid vector and the leaflet normal, with the lipid vector defined as the vector joining the center of mass of the terminal tail beads with the head phosphate bead.

## Results & Discussion

### Ripple formation in atomistic simulations of DPPC membranes

To robustly capture the ripple phase in all-atom DPPC membranes^26^, well-equilibrated, fluid, 368-lipid DPPC membranes at 350 K (time point I in Figure 1*a, c*) were cooled to 290 K, lower than the experimentally observed pre-transition temperature of 300-310 K^34,36–38^ and sufficient to ensure leaflet interdigitation^18,46^, using the stochastic simulated annealing MD algorithm, followed by constant temperature equilibration MD (EQMD) in the NPT ensemble at 290 K to ensure thermal equilibration and structural convergence (details in Methods; 2 independent MD replicates). Along with the initial state (time point I), three other time points are important in the combined SAMD+EQMD trajectory (Figure 1*a, c*) – the fluid membrane configuration where interdigitation starts (time point II), the fully interdigitated ripple phase membranes at the end of simulated cooling (time point III), and the converged systems at the end of constant temperature equilibration (time point IV). The extent of interdigitation was tracked by the minimum inter-leaflet P-P distance with time (ξ; Methods and Equation 1; Figure 1*b, c*). A higher initial value marked the baseline corresponding to the fluid phase (350 K; time point I), and an abruptly decreasing trend indicated significant interdigitation of lipids from opposite leaflets, with the onset of interdigitation marked by the inflection point (time point II – τ; see Methods, Figure S2, and Equation 1). By the end of simulated cooling (290 K; time point III), interdigitation was complete and a membrane with an ‘asymmetric’^20^ 1D ripple was formed, similar in structure to previous studies^19,20,45,46^, structurally stable upon further 200 ns equilibration at 290 K (time point IV), and with clearly distinct major and minor arms (illustrated in Figure S1*a*).

Voronoi-thickness membrane maps, which are Voronoi diagrams of the upper membrane leaflet superimposed onto P-P thickness heatmaps^25,43^, are shown in Figure 1*d*, corresponding to the top views of MD snapshots in Figure 1*a*. The trends were similar^43^ when the Voronoi diagrams were plotted using the lower leaflets (not shown). Compared to the uniform topography in the fluid phase, the formation of interdigitated (blue) and gel (yellow) domains in the ripple phase can be clearly observed. There is negligible difference in membrane topographies between time points III and IV except for the lateral movement of membrane across period boundaries, indicating that the ripple phase membrane configuration at the end of simulated cooling is at equilibrium. An interesting observation was that the interdigitated ripple domain (Figure 1*d*) appeared to be tilted in the lateral plane, whereas ripple domains in previous studies^19,20,43^ were typically parallel to one of the lateral box edges. Previous mesoscopic lipid bilayer simulations^21^ have revealed that at the highest value of the lipid head-head repulsion parameter explored, lower and higher values of which either abrogated or stabilized the ripple phase, respectively, the bilayer formed a laterally tilted ripple domain and a stable *P*_β′_ membrane phase similar to our observations.

Another independent all-atom MD replicate also yielded a similar interdigitated DPPC ripple phase membrane upon cooling (Figure S1*b*), thus validating the robustness of the above MD protocol to produce the *P*_β′_ phase from fluid membrane configurations. We next employed this MD cooling protocol to fluid-phase MARTINI membranes and evaluated their ability to capture the ripple phase.

### Ripple formation in smaller MARTINI membranes

Fluid phase, 380-lipid, DPPC MARTINI membranes belonging to six MARTINI FF variants were cooled from 350 K to 280 K, and evaluated for their ability to form the interdigitated *P*_β′_ membrane phase (see Introduction and Methods). The lower bound for temperature was set to 280 K, compared to 290 K for all-atom, because a previous CGMD study showed that ripple-like interdigitated phases in MARTINI DPPC membranes were formed between 280-290 K^36^. The trajectory of an SAMD+EQMD replicate with the MARTINI2n FF is shown with time points I-IV highlighted (Figure 2*a-c*). The constant temperature EQMD phase post cooling was chosen to be 5 μs for MARTINI (Figure 2*b*), compared to 200 ns for the all-atom simulations (Figure 1*c*). Similar to the time course seen in atomistic DPPC membranes, a majority (70%), but notably not all, MARTINI2n replicates showed interdigitation of membrane leaflets (time points I-IV on the ξ(*t*) plot, Voronoi-thickness maps pertaining to MD snapshots at time points I and IV), thus apparently capturing ripple-like membrane states. This is in accord with previous single replicate studies, where MARTINI membranes of similar sizes captured the ripple phase^31,36,48,49^. In the rest of the MARTINI2n replicates (30%), the membranes bypassed the interdigitated ripple-like state and directly froze into *L*_β_ configurations (Figure S2). This emphasizes that with simulated cooling MD, rippling in MARTINI membranes depended on the initial configuration and starting velocities, and therefore necessitated multiple MD replicates for obtaining robust trends.^52^

MARTINI SAMD simulations showed two differences compared to all-atom SAMD simulations. Firstly, during SAMD, the atomistic DPPC membranes stay in the fluid phase with disordered tails before the onset of interdigitation (Time point II in Figure 1*a*) and therefore *m*_l_ ≈ 0 (Figure 1*c*; Equation 1). In contrast, *m*_l_ > 0 for the MARTINI DPPC membranes since they exhibit a degree of tail ordering and expansion along the Z-axis (compensated by reduced X-Y dimensions) with decreasing temperature before the start of interdigitation (Time point II in Figure 2*a*), compared to the fluid phase at 350 K (Time point I in Figure 2*a*). Secondly, the orientation of the interdigitated domains in the MARTINI2n membranes were perpendicular to the x-axis (Figure 2*c*), in contrast with the laterally tilted interdigitated domains seen in all-atom simulations (Figures 1*d* and S1*b*). Similar domains were observed in a previous MARTINI study.^36^

Formation of ripple-like phases in DPPC membranes described by the other FF variants – MARTINI2On, MARTINI3a, MARTINI3b, MARTINI2p, and MARTINI2Op – are summarized in Figures 2*d-g* and S5*b* (all MD replicates for each FF variant displayed in Figures S3-S7). MARTINI2On is similar to MARTINI2n, with 70% of the replicates forming ripple-like (Figures 2*d* and S3). MARTINI3a, the improved^33,51^ next generation MARTINI3 FF, showed ripple-like formation in 70% of the replicates, similar to MARTINI2n and MARTINI2On (Figures 2*e* and S4). (The beta version of MARTINI 3 – MARTINI3b – showed ripple-like formation in 20% of the replicates (Figure S5).) The MARTINI FF combinations with polarizable water, MARTINI2p (Figures 2*f* and S6) and MARTINI2Op (Figures 2*g* and S7), showed ripple-like states in only 10% and 30% of replicates, respectively, despite the reparameterization of polarizable water to improve the accuracy of solvent electrostatic effects^50^, important for capturing macroscopic membrane properties^57,58^. Our observations with MARTINI2p are to be contrasted with previous single-replicate MARTINI^36^ simulations, where quasi-static cooling of fluid-phase membranes resulted in the formation of the ripple-like domains only with polarizable water. While differences in MD protocols and differences in the initial configurations may explain the disparities, careful future investigations, with systems of varying size, may be necessary to determine the predictive accuracy of polarizable water-lipid interactions.

### Testing for ripple formation in large atomistic and MARTINI DPPC membranes upon simulated cooling

In a previous study, Rodgers *et al.*^36^ observed the interdigitation of lipids in relatively small 512-lipid MARTINI DPPC membranes upon quasi-equilibrium, stepwise, simulated cooling of fluid membranes from 325 K to less than 280 K. These small membranes apparently sampled an interdigitated ripple-like phase, similar to our observations (Figure 2). However, when the exercise was repeated with 4-times-larger (2048-lipid) DPPC membranes, the ripple-like phase could not be reproduced. Thus, Rodgers *et al.*^36^ concluded that the interdigitated ripple-like states in smaller membranes were actually ‘intermediate’ kinetically trapped membrane states rather than a true ripple phase. In contrast to these MARTINI simulations, a large all-atom 3200-lipid DPPC membrane has been shown to form an interdigitated ripple phase^31^, albeit with a different MD protocol. Therefore, we next tested ripple formation in large all-atom DPPC membranes with our simulated cooling MD protocol (similar to Figure 1), that would re-establish a benchmark to test for ripple formation and system size effects in large MARTINI membranes.

We set up two 2208-lipid DPPC bilayer configurations, which were 6-times-larger than the smaller atomistic DPPC membranes shown in Figure 1, and larger than the membranes employed by Rodgers *et al.*^36^, and pre-equilibrated them in the *L*_α_ phase at 350 K (Figure 3*a*). Subsequently, these membranes were cooled to 290 K with the same SAMD protocol. One of the all-atom MD replicates in the fluid phase formed an interdigitated ripple phase membrane by 290 K (Figure 3*b*). The other replicate (Figure S8) formed a non-interdigitated ‘symmetric’ ripple^20^ in the larger dimension, which is typically observed upon slow cooling from the fluid phase to approximately the gel-to-fluid transition temperature^20^, and an asymmetric interdigitated ripple (similar to Figures 1, 3) in the smaller dimension. Therefore, both smaller and larger all-atom DPPC membranes, described by the CHARMM36 FF, sample the thermodynamic ripple phase upon simulated cooling MD without any evidence of finite size artefacts. In contrast to smaller membranes (Figures 1*d*, S9a, and S1), the Voronoi-thickness map of the first large membrane replicate shows reticulated ripple domains (Figures 3*b* and S9*b*). Our observations are in accord with previous studies^31,45^, where simulated quenching of 1280-lipid and 3200-lipid all-atom DPPC membranes to below the main transition temperature also resulted in reticulated ripple domains. Significantly longer runs are probably required to anneal and reorganize these reticulated ripple domains into classically observed periodic ripple domains.

**Figure 3.**
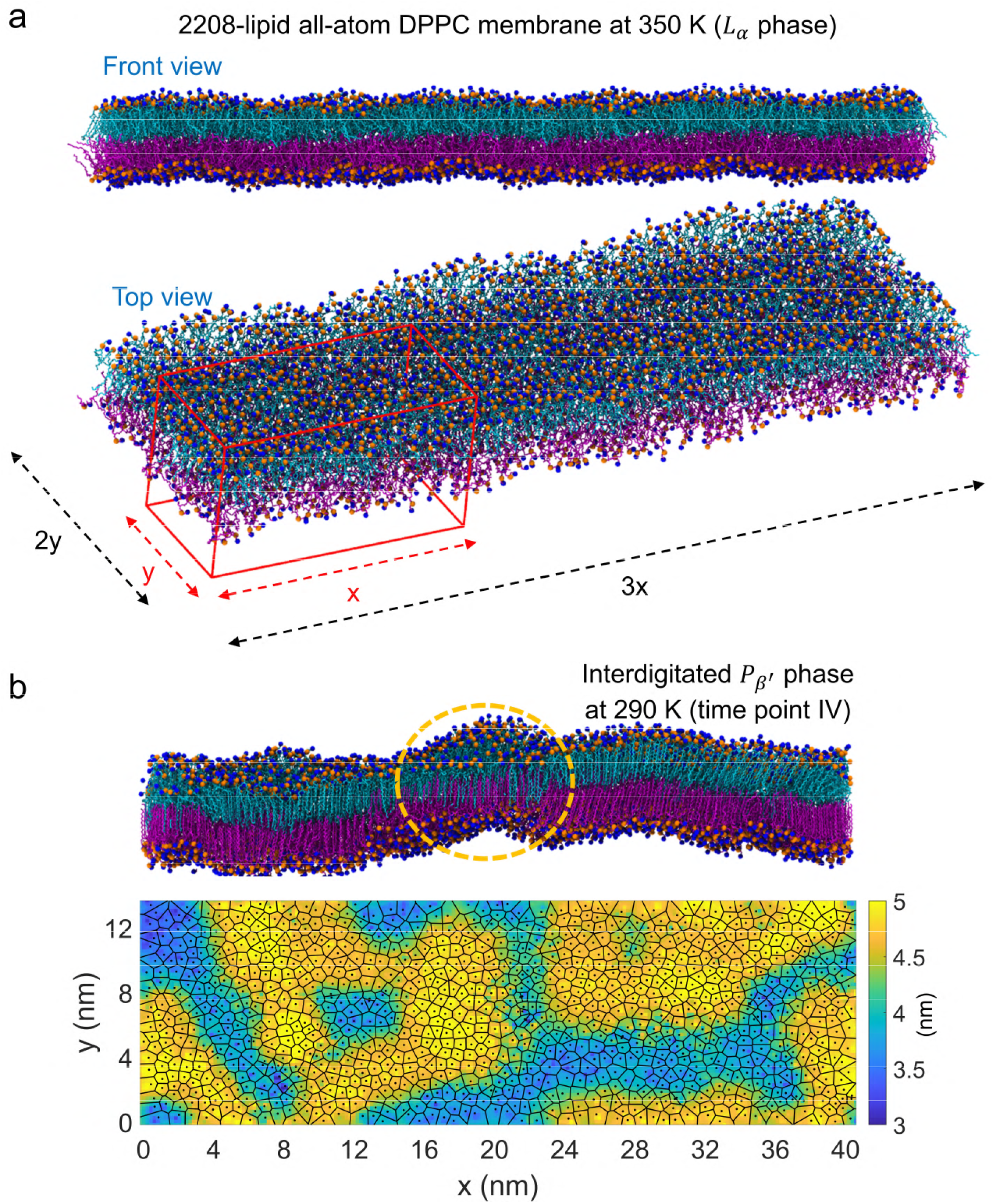
Ripple formation from all-atom MD of a large DPPC membrane. **(a)** Front and top views of a large 2208-lipid DPPC membrane which was pre-equilibrated in the fluid phase at 350 K. For comparison, a representative simulation box of a smaller all-atom membrane (Figure 1) is shown in red. The large membrane is 6-times (3X times 2Y) larger than the smaller membrane. **(b)** Front view of the large membrane after cooling to 290 K and EQMD (corresponds to time point IV), and its Voronoi-thickness modulation map (blue represents interdigitated regions) shows the formation of an aperiodic and reticulated interdigitated ripple domain (also see Figures S8-S9).

We next tested whether large, fluid, rectangular, 2204-lipid MARTINI DPPC membranes, with normal (not polarizable) MARTINI water, also sampled the interdigitated ripple-like phase upon cooling from 350 K to 280 K. Two initial configurations each for MARTINI2n, MARTINI2On, MARTINI3a, and MARTINI3b FFs were set up analogous to the large all-atom membranes in Figure 3, and pre-equilibrated at 350 K, followed by cooling to 280 K (Figures 4 and S10). However, none of the large MARTINI membranes captured a ripple-like phase upon cooling to 280 K. MARTINI2n and MARTINI3a/b membranes directly froze into the *L*_β_ phase, while the MARTINI2On membranes transitioned to the *L*_β′_ phase (Figure 4b); confirmed by independent MD replicates (Figure S10). To ensure that membrane geometry did not influence these results, we repeated these simulations with large 6728-lipid square MARTINI2On membranes, and these also did not sample ripple-like states (Figure S11). Since only smaller (380-lipid) but not larger (>2204-lipid) DPPC MARTINI membranes captured the interdigitated ripple-like states, in contrast to all-atom membranes, we inferred that the smaller MARTINI ripple-like membranes (Figures 2, S2-S7) represent kinetically-trapped states caused by finite box-size effects rather than true ripple states.

**Figure 4.**
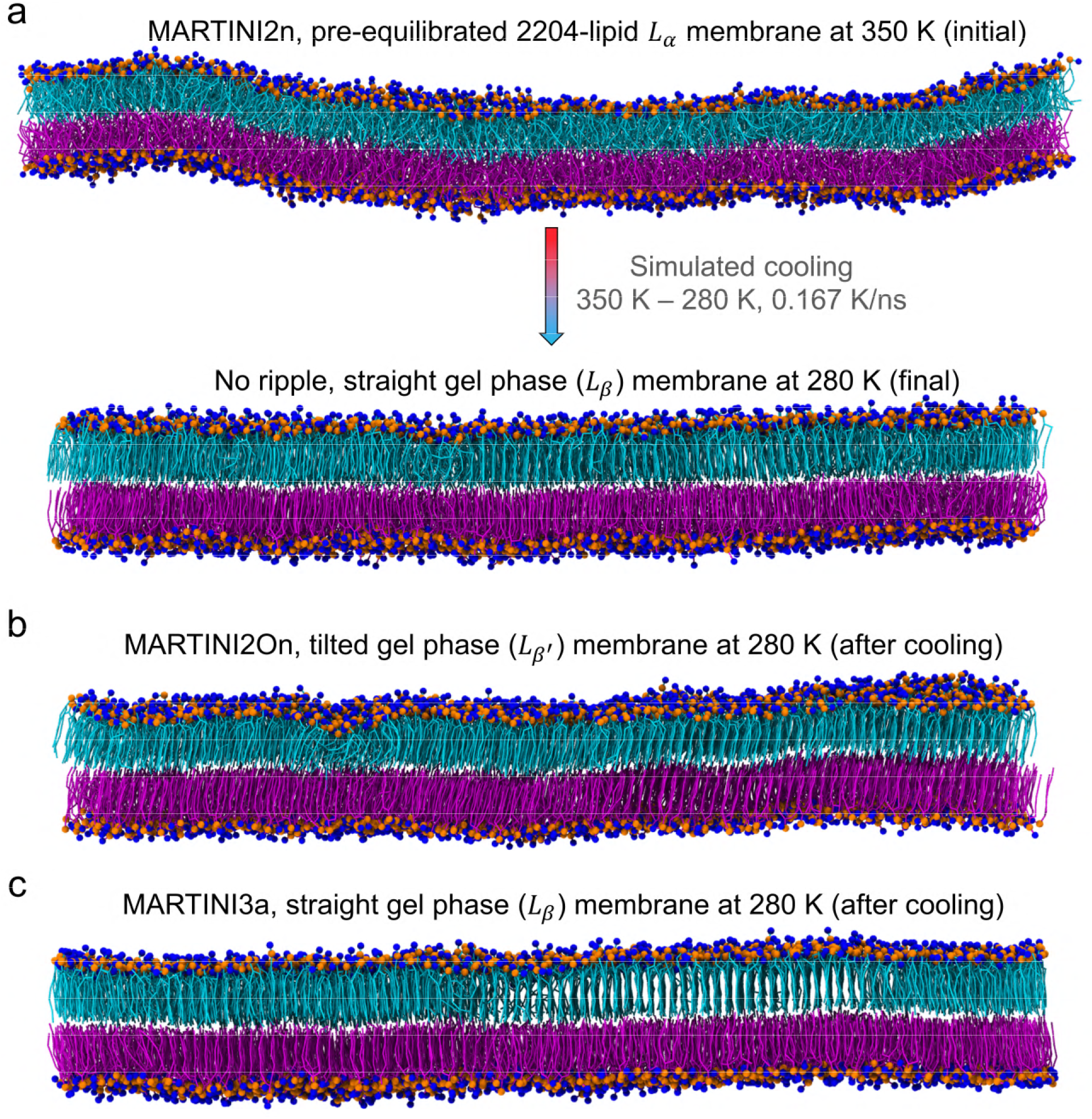
Lack of ripple formation in large MARTINI DPPC membranes. **(a)** A representative 2204-lipid, rectangular, MARTINI2n DPPC membrane, pre-equilibrated at 350 K in the *L*_α_ phase, was cooled directly into the gel phase at 280 K without the membrane sampling a ripple-like phase, unlike all-atom systems (see Figure 3). This behaviour was also observed with **(b)** MARTINI2On and **(c)** MARTINI3a FFs. Independent MD replicates are shown in Figure S10. Two independent simulation replicates, where large 6728-lipid, square, MARTINI2On membranes were cooled to 290 K, also showed a similar lack of sampling of a ripple-like phase (Figure S11), suggesting that system geometry did not influence our observations.

### Equilibrium MD simulations of large MARTINI membranes initialized in the ripple-like phase show loss of interdigitation

To confirm the aforementioned inference, we employed an orthogonal approach where multi-μs equilibrium simulations of large MARTINI membranes, initialized in periodic interdigitated ripple-like configurations, were performed at 280 K. These large systems were created from the final, interdigitated, ripple-like, smaller (380-lipid) MARTINI membrane configurations after 1 μs of constant temperature EQMD at 280 K. The smaller membranes were repeated 6-times along the lateral bilayer plane across periodic boundaries (3x times 2y), and the initial large 2280-lipid ripple-like membrane configurations thus created are displayed in Figures 5*a, c, d* and S12*d*, corresponding to MARTINI2On, MARTINI2n, MARTINI3a, and MARTINI3b, respectively.

**Figure 5.**
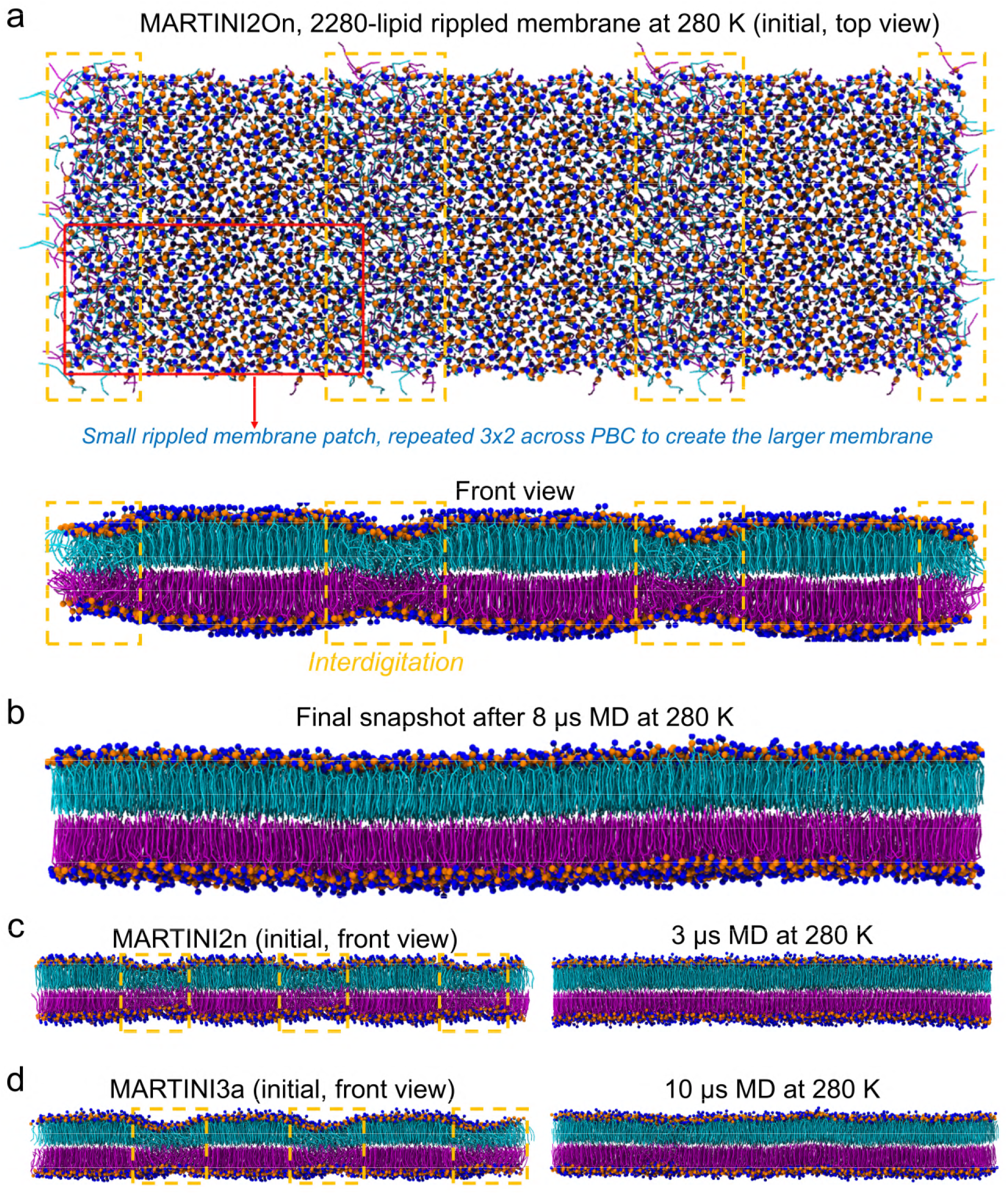
MD simulations of large MARTINI DPPC membranes starting with interdigitated ripple-like states. **(a)** To test the stability of kinetically trapped domains (yellow dashed rectangles) at larger system sizes, a 2280-lipid MARTINI2On DPPC membrane configuration with periodic ripples (top and front views) was created by repeating smaller ripple-like membranes (from time point IV, Figure 2d) six-times across periodic boundary conditions (thrice in X, twice in Y). **(b)**8 μs MD at 280 K almost completely abolished the interdigitated ripple-like configuration and led to the establishment of the *L*_β_ phase (Voronoi-thickness maps in Figure S12*a*). Similar loss of the ripple-like domains was seen for large MARTINI2n **(c),** and MARTINI3a **(d),** membranes (Voronoi-thickness maps in Figure S12*b, c*).

The rationale behind performing equilibrium MD of these large systems was simply to test if the ripple-like states in small systems were stabilized due to finite size effects. Upon 3-10 μs of constant temperature NPT-MD at 280 K, we observed that the large, interdigitated MARTINI membranes spontaneously froze into the gel phase, as shown by both MD snapshots (Figures 5 and S12*d*) and Voronoi-thickness maps (Figure S12*a-d*), indicating that the initial ripple-like membrane configuration was unfavourable at larger system sizes. This phenomenon was observed for all the four MARTINI FF variants, although the time scale over which interdigitation disappeared leading to formation of the final gel phase configuration varied amongst the FFs. MARTINI2On showed progressive loss of interdigitation which was nearly complete by 8 μs (Figures 5*b* and S10*a*). Longer simulations may have shown a complete loss of interdigitation, but we did not further pursue these computationally intensive MD simulations since the progressive loss of interdigitation with time was apparent (Figures 5*a, b* and S12*a*). MARTINI2n showed a rapid and complete loss of interdigitation by 3 μs (Figures 5*c* and S12*b*), and MARTINI3a/b showed near-complete loss of interdigitation by 10 μs (Figures 5*d* and S12*c, d*), similar to MARTINI2On. In contrast, none of the smaller membranes showed any loss of interdigitation upon 5 μs of constant temperature EQMD at 280 K (Figures 2 and S5*b*).

Together with observations from the simulated cooling of large fluid membranes, these two orthogonal MD approaches (Figures 4, 5, S10-12) strongly suggest that MARTINI membranes (with all the tested FFs) do not capture the thermodynamic ripple phase, unlike all-atom membranes. Our systematic study implies that ripple-like phase formation observed in smaller MARTINI membranes was due to kinetic trapping of interdigitated leaflets at small system sizes. The few studies which indeed report MARTINI ripple phase membranes, both with^49^ and without^36,48^ enhanced sampling, are with smaller membranes composed of 390^49^ and 512^36,48^ lipids, respectively. Therefore, based on our observations, the formation of rippled bilayers in these studies could be attributed to finite size artefacts^36^ until they are validated with simulations at larger system sizes. Thus, MARTINI FF parameters may have to be tuned to better reproduce the lipid membrane phase diagram, and the *P*_β′_ phase in particular. FF approaches that tune the bonded parameters of MARTINI lipids based on comparison with all-atom simulations^25,31,36^ may result in models that better capture membrane phase behaviour.

### The optimized MARTINI DPPC FF (MARTINI2On) forms the *L*_β′_, but not *P*_β′_, phase

DPPC, with its large headgroup and strong head-head interactions, favours a tilted gel (*L*_β′_) phase at lower temperatures. Therefore, during the *L*_α_ →; *L*_β′_ transition upon simulated cooling, the tilted ripple (*P*_β′_) state occurs. Lipids exhibiting an *L*_β_ state at lower temperatures, such as phosphatidylethanolamines^75^, show a single phase transition and do not form the ripple phase^20^. In our small membrane simulations, MARTINI2n (Figure S2) and MARTINI3a/b (Figures S4 and S5) mostly froze into *L*_β_ states below transition temperatures, in accord with previous studies^36,49^. MARTINI2n was previously shown to sample the *L*_β′_ phase only upon enhanced sampling^76^. The frozen lipids in the upper leaflet of replicate 2 of MARTINI3a (Figure S4) are weakly tilted, suggesting that this FF may also capture the *L*_β′_ phase upon enhanced sampling. This propensity to freeze into the *L*_β_ state may underlie the inability of these FFs to capture the *P*_β′_ phase. However, MARTINI2On, possessing optimized DPPC bond parameters that closely match macroscopic membrane properties from all-atom simulations, spontaneously captured the *L*_β′_ phase in many of the small membrane replicates upon cooling with classical MD simulations (Figure S3). This agrees with a previous study^31^ that developed the MARTINI2On DPPC parameters, where the authors also observed the formation of the *L*_β_1 phase without enhanced sampling. Even in replicates that formed interdigitated ripple-like configurations (Figure S3), the non-interdigitated gel-like domains often resembled the *L*_β_1 phase.

To further contrast the lipid tilt in the gel phase configurations of DPPC MARTINI2n and MARTINI2On membranes, we chose representative replicates (#7 in Figure S2, #3 and #7 in Figure S3) that had frozen into gel states after SAMD without sampling interdigitated ripple-like states, and further equilibrated them at 280 K for 5 μs in the NPT ensemble. The final MD snapshots for MARTINI2n (#7 from Figure S2) and MARTINI2On (#7 from Figure S3) are illustrated in Figure 6*a, b*. Clearly, MARTINI2n and MARTINI2On form stable *L*_β_ and *L*_β′_ configurations, respectively, at 280 K. For both systems, we quantified the extent of lipid tilt from the tilt angle (θ°) distributions of lipids with their respective leaflet normal (Methods; Figure 6*c*). The MARTINI2n and MARTINI2On gel phase tilt distributions were observed to have modes at 7.3° and 15.2°, respectively.

**Figure 6.**
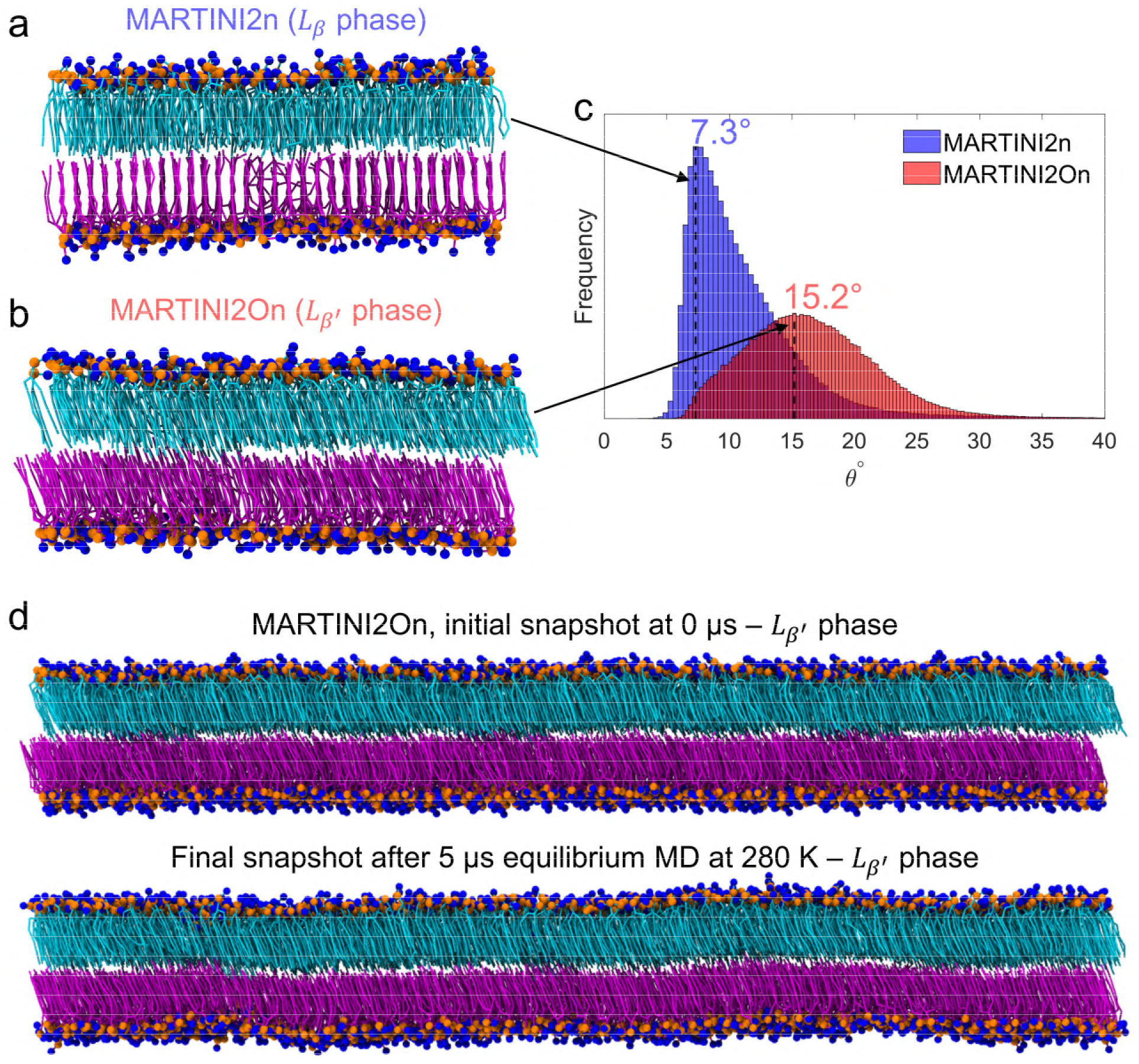
Comparison of gel-phase structures of MARTINI membranes. Gel phase DPPC bilayer structures with **(a)** the standard MARTINI2n FF, and **(b)** the optimized MARTINI2On FF are shown. MARTINI2n membranes freeze into the straight gel phase (*L*_β_) whereas MARTINI2On membranes often sample the tilted gel phase (*L*_β′_), quantified by the lipid tilt-angle distributions **(c)**. **(d)** To confirm that this was a true thermodynamic phase and not a finite size effect, the smaller *L*_β′_ membrane in (a) was replicated 6x (initial snapshot) and then equilibrated for 5 μs at 280 K. The membrane remained in the *L*_β′_ phase thus ruling out finite size artefacts.

To ensure that the MARTINI2On *L*_β′_ configuration was not a consequence of finite size artefacts, we created a 6-times-larger membrane by repeating the membrane from Figure 6*b* across periodic boundaries (3x times 2y; similar to Figure 5). This initial configuration is shown in Figure 6*d*, and was simulated at 280 K for 5 μs. From the final snapshot (Figure 6*d*), the *L*_β′_ configuration is completely preserved. Additionally, simulated cooling of a large *L*_α_ MARTINI2On membrane from 350 K – 280 K also resulted in a *L*_β′_ configuration (Figure 4*b*). These observations strongly suggest that MARTINI2On membranes can sample the true *L*_β′_ phase at lower temperatures, without any finite size artefacts, and in agreement with previous studies^31^. However, our multi-replicate simulations show that MARTINI2On membranes also stochastically sample the *L*_β_ phase, as shown by replicates #2 and #8 in smaller membranes (Figure S3), and from the simulations of large membranes (Figures 5*b*, S10 and S11). Therefore, there may be room for further optimization of the MARTINI2On DPPC FF for accurate reproduction of low temperature phase behaviour.

Interestingly, when replicate #3 from Figure S3 was equilibrated for 5 μs at 280 K, it showed the formation of a weakly interdigitated titled domain (Figure S13*a*), a potential precursor for the formation of a true ripple phase. To assess this further, we set up a 6-times larger membrane (Figure S13*b*) by repeating the smaller system across periodic boundaries along the membrane plane (similar to Figure 5*a*). After 10 μs of constant temperature EQMD at 280 K, the interdigitation almost completely disappeared as shown by the final MD snapshot (Figure S13*b*) and corresponding Voronoi-thickness maps (Figure S13*c*).

## Conclusions

Molecular simulations at various levels of FF descriptions, ranging from all-atom and united atom models at the finer scales to DPD and MARTINI models at higher levels of coarse graining, have been extensively used to study the rich variety of equilibrium phases observed in model phospholipid bilayer membranes^1,17,77^. In particular, the rippled *P*_β′_ phase, sensitive to initial configurations, annealing protocols, and finite-size effects, has been challenging to capture using MD simulations^20,25,36,43^. Although all-atom FFs have successfully captured the formation of the ripple phase (Table 1), the ability of the widely used MARTINI force-fields to stabilize this intermediate phase, which occurs below the main gel-to-liquid crystalline transition, has been a matter of some debate^36,49^.

In this manuscript, we assessed the ability of MARTINI DPPC lipid membranes described by six MARTINI FF variants (see Introduction and Methods) towards capturing the ripple phase. Previous experiments^35,39–42^ and all-atom simulations^19,31,45,46^ have shown that DPPC membranes sample the tilted ripple phase at pre-transition temperatures. Previous MARTINI studies yielded conflicting observations, with some inferring that MARTINI DPPC lipid membranes could not capture the ripple state^36^, while other studies observed otherwise^31,48,49^. Therefore, we established a standard simulated annealing-based MD protocol, built on insights from previous studies^18–20,31,45,46^, for reliably producing the tilted ripple phase in multiple all-atom DPPC membrane MD replicates. By cooling from the fluid phase to below the phase transition followed by constant temperature equilibration. The structure of these rippled membranes resembled those from previous studies^19,20,45,46^. We then applied the same MD protocol to fluid phase MARTINI DPPC membranes with six MARTINI FF variants and ten independent MD replicates per variant. Simulations with all the MARTINI FFs showed that a fraction of replicates formed interdigitated ripple-like states. These simulations also reinforced the importance of using multiple independent MD replicates for robustly observing phase behaviour^18,52^. The lateral orientation of interdigitated domains in these ripple-like membranes were different compared to those from all-atom simulations and resembled a membrane with co-existing interdigitated *L*_α_ and gel phases.

Next, we explored whether the MARTINI ripple-like membranes were kinetically trapped states enforced by finite system size effects. To this end, we first re-established the all-atom protocols by employing the aforementioned MD cooling protocol on large all-atom DPPC membranes. We observed that these larger membranes also formed the tilted ripple phase with reticulated interdigitated domains, in agreement with previously reported large-membrane all-atom DPPC simulations^31,45^. However, when similarly large MARTINI DPPC membranes (4 FF variants, 2 replicates each) were cooled from the fluid phase, they failed to capture ripple-like states and froze directly into the gel phase. To confirm this behaviour via an orthogonal approach, we created larger membranes from previously formed smaller ripple-like membranes, by repeating them across periodic boundaries. These larger MARTINI periodic ripple-like membranes (4 FF variants), with an initial configuration bias towards the ripple phase, were then simulated at a constant temperature of 280 K, below the transition temperature. Strikingly, all the membranes showed a progressive loss of the interdigitated ripple-like domains, and either transitioned, or were in the process of transitioning, into the gel phase. Based on these findings we attribute the ripple-like states observed in the smaller MARTINI membranes to finite size artefacts. In the case of the MARTINI2On DPPC FF, the inability to form the tilted ripple was despite its ability to capture the tilted gel phase at lower temperatures without the need for enhanced sampling.

In summary, our systematic observations suggest that the current MARTINI DPPC FFs, including the recently developed MARTINI3, may not be able to capture the *P*_β′_ phase, which involves an intricate interplay of lipid tail interdigitation, tilt and headgroup hydration in the ripple state of the membrane. CG force fields like MARTINI are known to overestimate enthalpy to compensate for an intrinsically low entropy due to reduced degrees of freedom^78^. Specifically reparametrizing the MARTINI FFs to correct this enthalpy-entropy imbalance and better reproduce membrane thermodynamics as well as phase behaviour is expected to yield improved CG models that may bridge the qualitative and quantitative gap towards all-atom MD simulations, while unlocking the speed and scaling advantages associated with the versatile and powerful MARTINI framework.

## Acknowledgements

We thank the Supercomputer Education and Research Center (SERC) computational facility at the Indian Institute of Science, Bengaluru, for access to supercomputing resources. K.G.A. acknowledges funding by a grant from the Department of Science and Technology, Government of India.

## Supporting information

Supporting information contains detailed author contributions, and thirteen supporting figures, namely: an all-atom MD simulation replicate reproducing the ripple phase (Figure S1), ten independent cooling MD simulation replicates with the six MARTINI FFs (Figures S2-S7), and large all-atom (Figures S8 and S9) and MARTINI (Figures S10-S13) membrane simulations.

## Supporting information

### Author contributions

Conceptualization: R.D.; Data curation: P.S.; Formal analysis: R.D. and P.S.; Funding acquisition: K.G.A.; Investigation: P.S. and R.D.; Methodology: R.D. and P.S.; Project administration: P.S. and R.D.; Resources: P.S., R.D. and K.G.A.; Software (simulations, analysis code): P.S. and R.D.; Supervision: K.G.A.; Validation: P.S., R.D. and K.G.A.; Visualization: P.S. and R.D.; Writing – original draft: R.D. and P.S.; Writing – review & editing: R.D., P.S. and K.G.A.

All authors have given approval to the final version of the manuscript.

### Supporting figures

**Figure S1.**
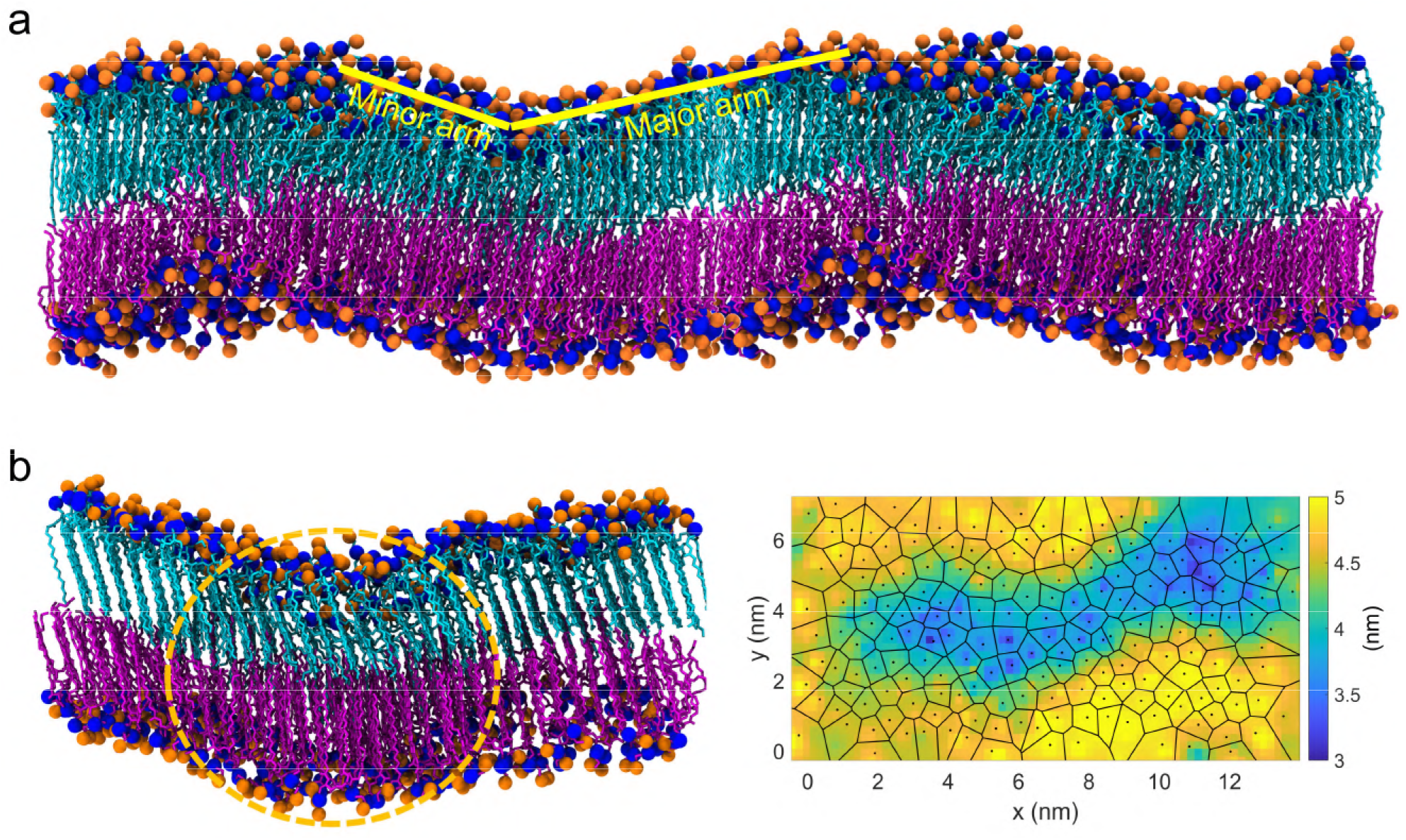
An independent all-atom MD replicate reproduces the ripple phase. **(a)** MD snapshot at time point IV from Figure 1*a* is periodically repeated along the x-axis, and the major and minor arms of the ripple in the upper leaflet are visually annotated for clarity. **(b)** Final snapshot (time point IV in Figure 1) and corresponding Voronoi-thickness diagram from the independent simulated cooling replicate, using the same MD protocol as for the replicate in Figure 1, shows robust reproduction of the tilted ripple phase from all-atom simulations.

**Figure S2.**
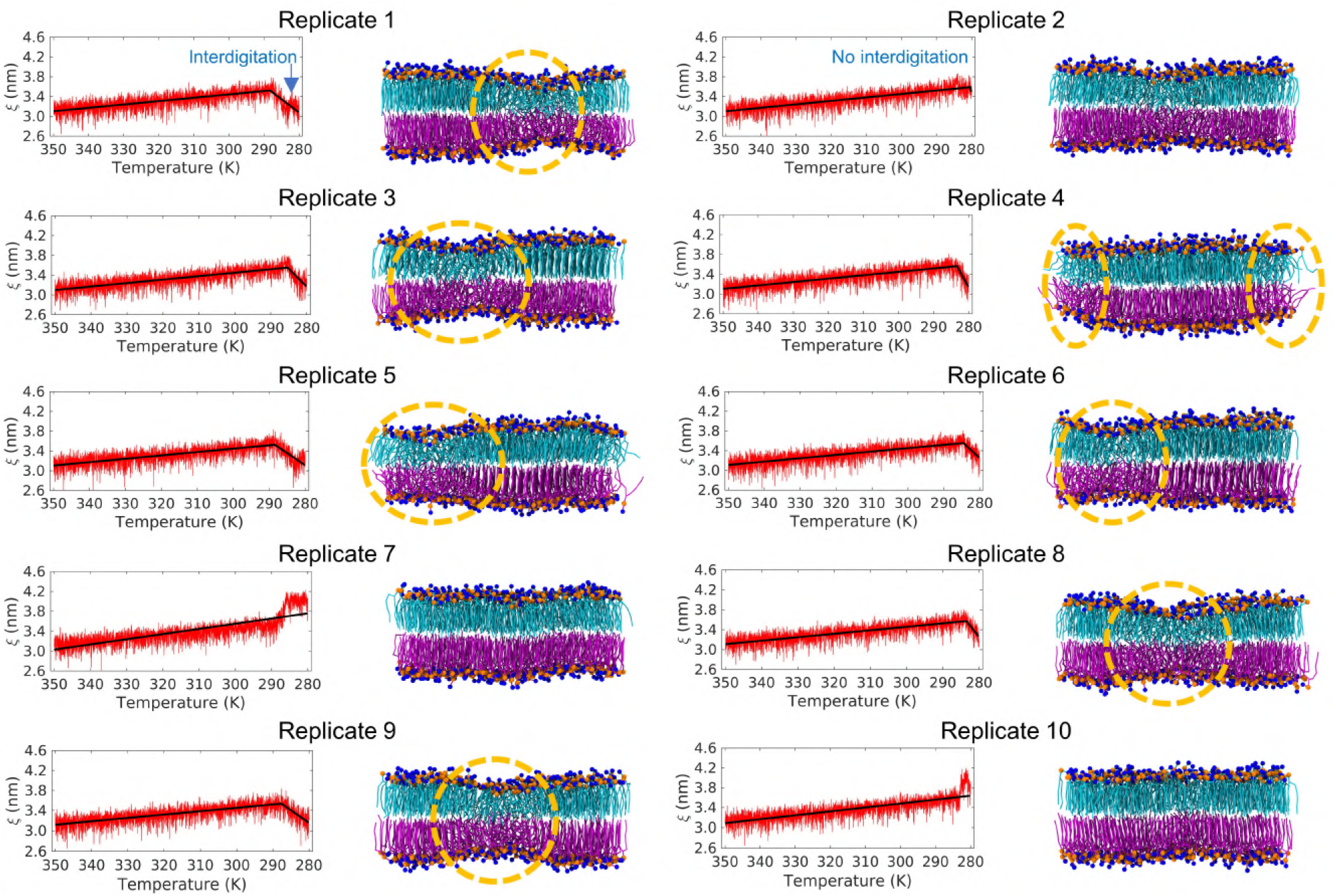
Summary of 10 independent cooling MD replicates with the MARTINI2n FF. Temporal evolution of the minimum inter-leaflet P-P distance (ξ), a measure of the extent of interdigitation (Figure 1*b*; *m*_2_ < 0 – interdigitation, and *m*_2_ > 0 – no interdigitation (Methods, Eq. 1), are highlighted in replicates 1 and 2, respectively). Final snapshots corresponding to time point III at the end of cooling simulations with MARTINI2n DPPC membranes are shown for all the 10 independent MD replicates. The interdigitated domains in replicates which exhibit ripple-like structures are marked with yellow dashed circles.

**Figure S3.**
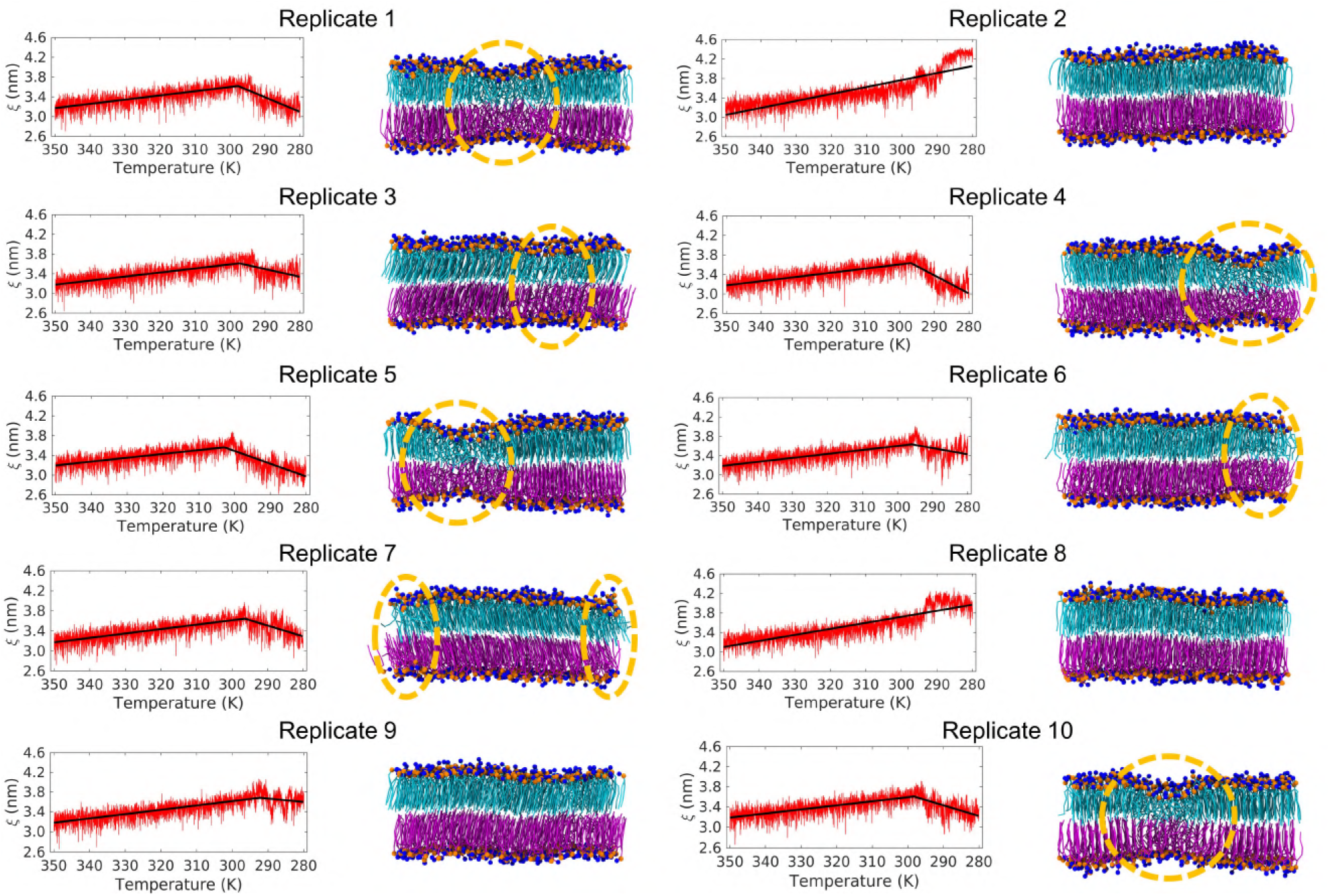
Summary of 10 independent cooling MD replicates with the MARTINI2On FF. Same as Figure S2, but with the MARTINI2On FF variant.

**Figure S4.**
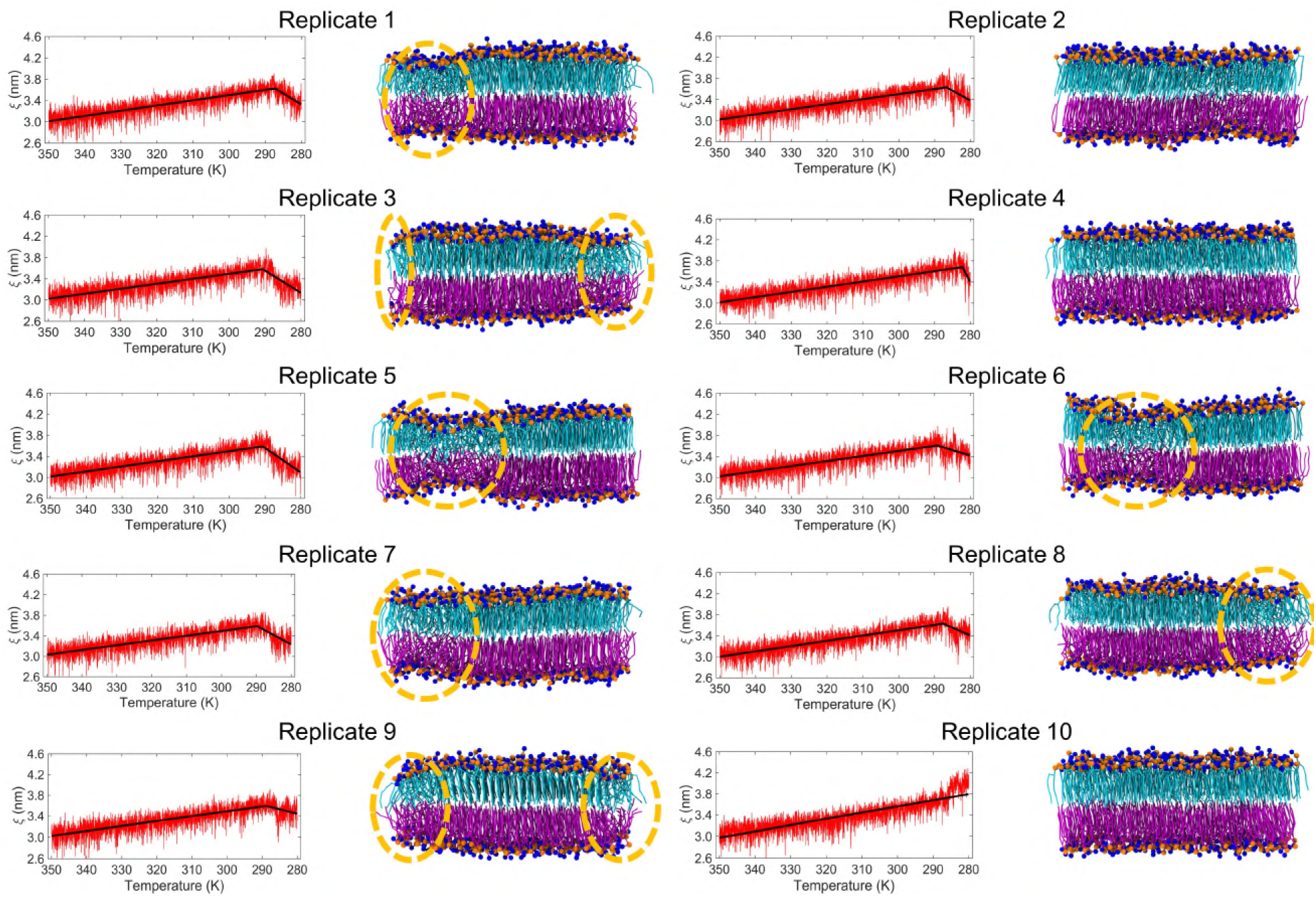
Summary of 10 independent cooling MD replicates with the MARTINI3a FF. Same as Figure S2, but with the MARTINI3a FF variant. Note that while the ξ trends for replicates 2 and 4 appear to indicate the onset of ripple formation, it can be inferred from visual inspection that this is due to very few interdigitating lipids and not due to the formation of a ripple-like domain. Hence, these replicates were not counted as ripple-like states.

**Figure S5.**
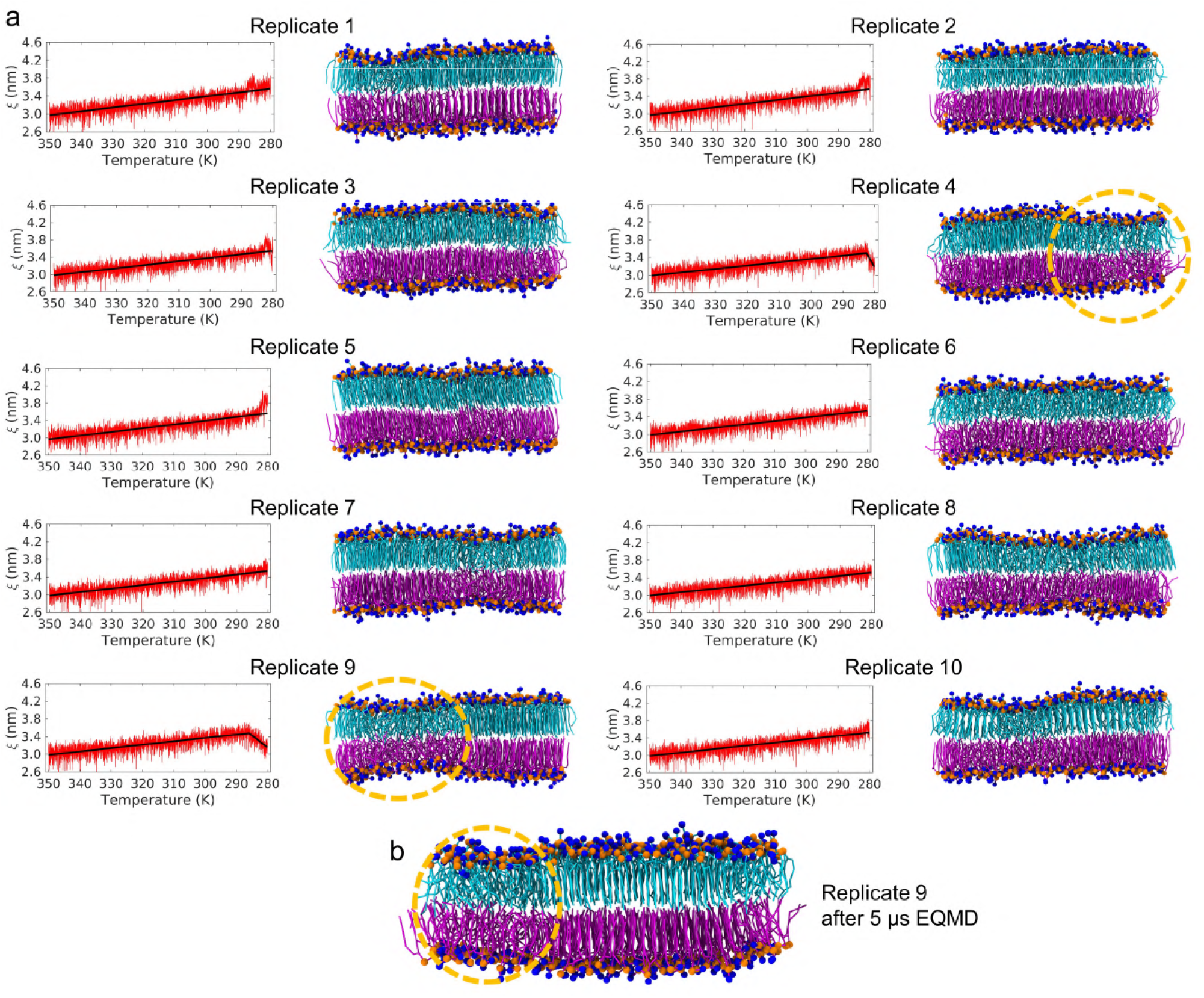
Summary of 10 independent cooling MD replicates with the MARTINI3b FF. **(a)** Same as Figure S2, but with the MARTINI3b FF variant. **(b)** Replicate 9 was further subjected to 5 μs EQMD to confirm that the ripple-like domains were stable at this timescale.

**Figure S6.**
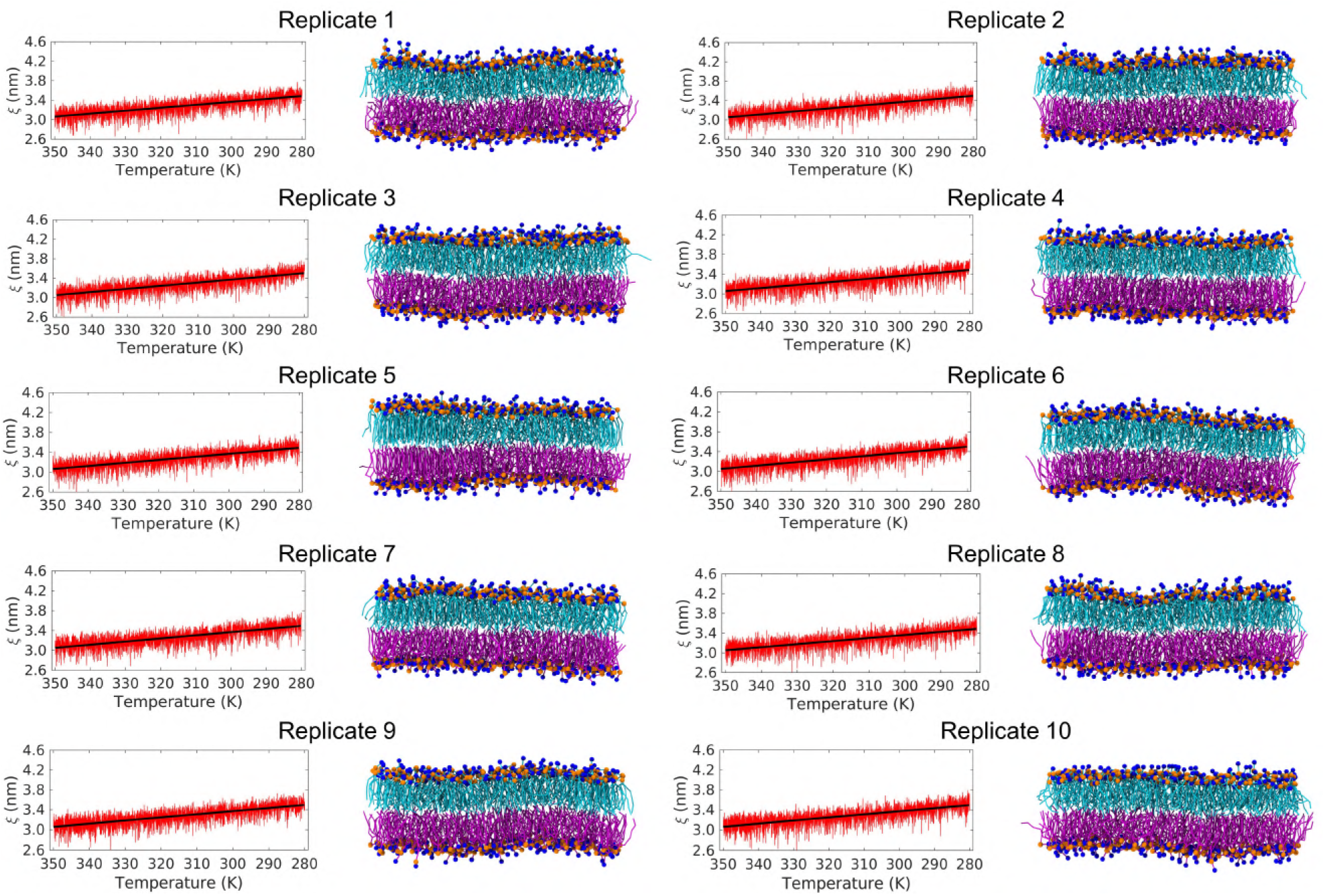
Summary of 10 independent cooling MD replicates with the MARTINI2p FF. Same as Figure S2, but with the MARTINI2p FF variant. Note that while none of the *ξ* plots show ripple-like domain formation during the cooling phase, replicate 5 formed shallow interdigitated ripple-like domains during the 5 μs EQMD (Figure 2*f*).

**Figure S7.**
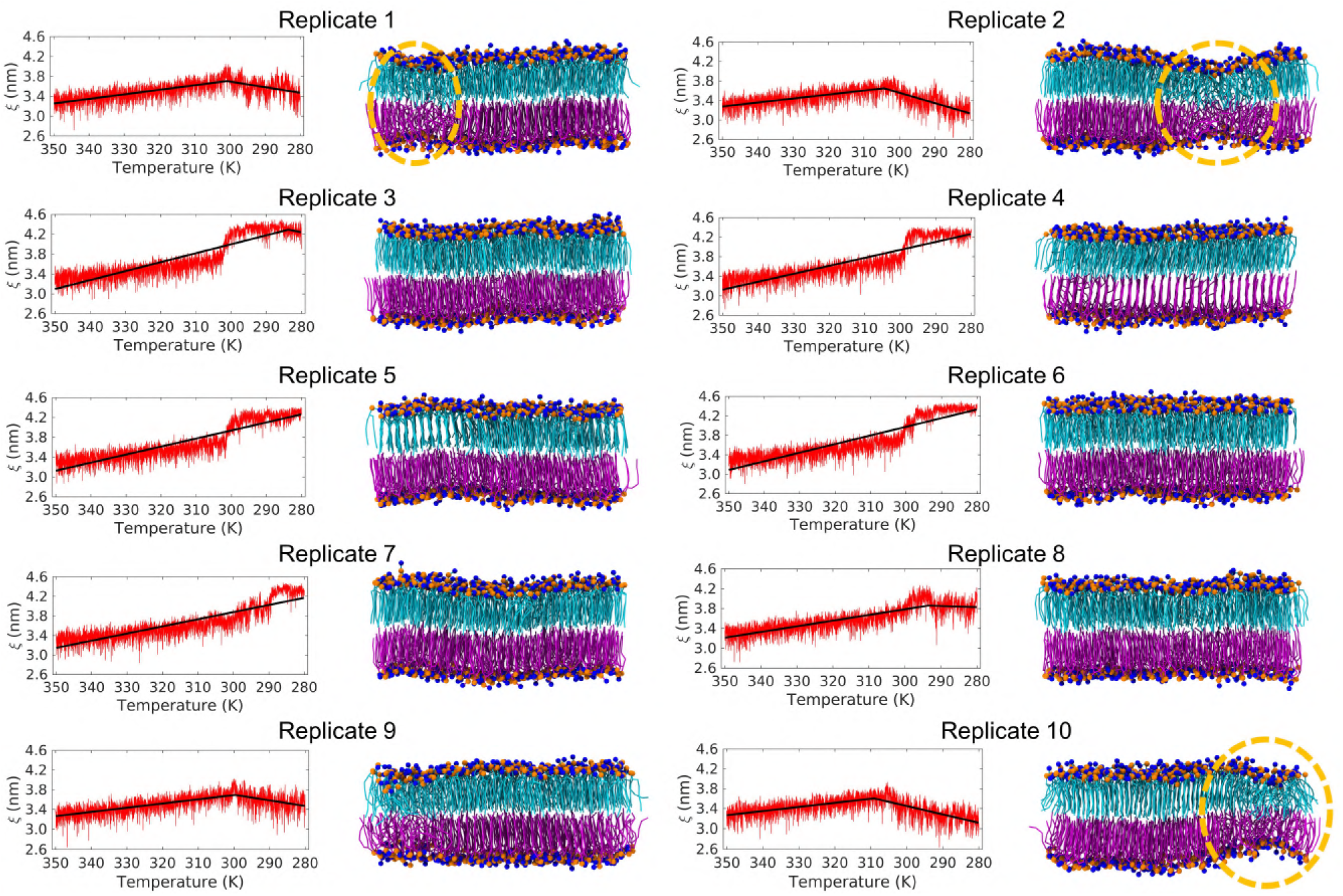
Summary of 10 independent cooling MD replicates with the MARTINI2Op FF. Same as Figure S2, but with the MARTINI2Op FF variant. In replicate 9, while we observed a ξ trend similar to ripple-like domain formation, visual inspection yielded no interdigitated domains, and hence this replicate was not considered as a ripple-like state.

**Figure S8.**
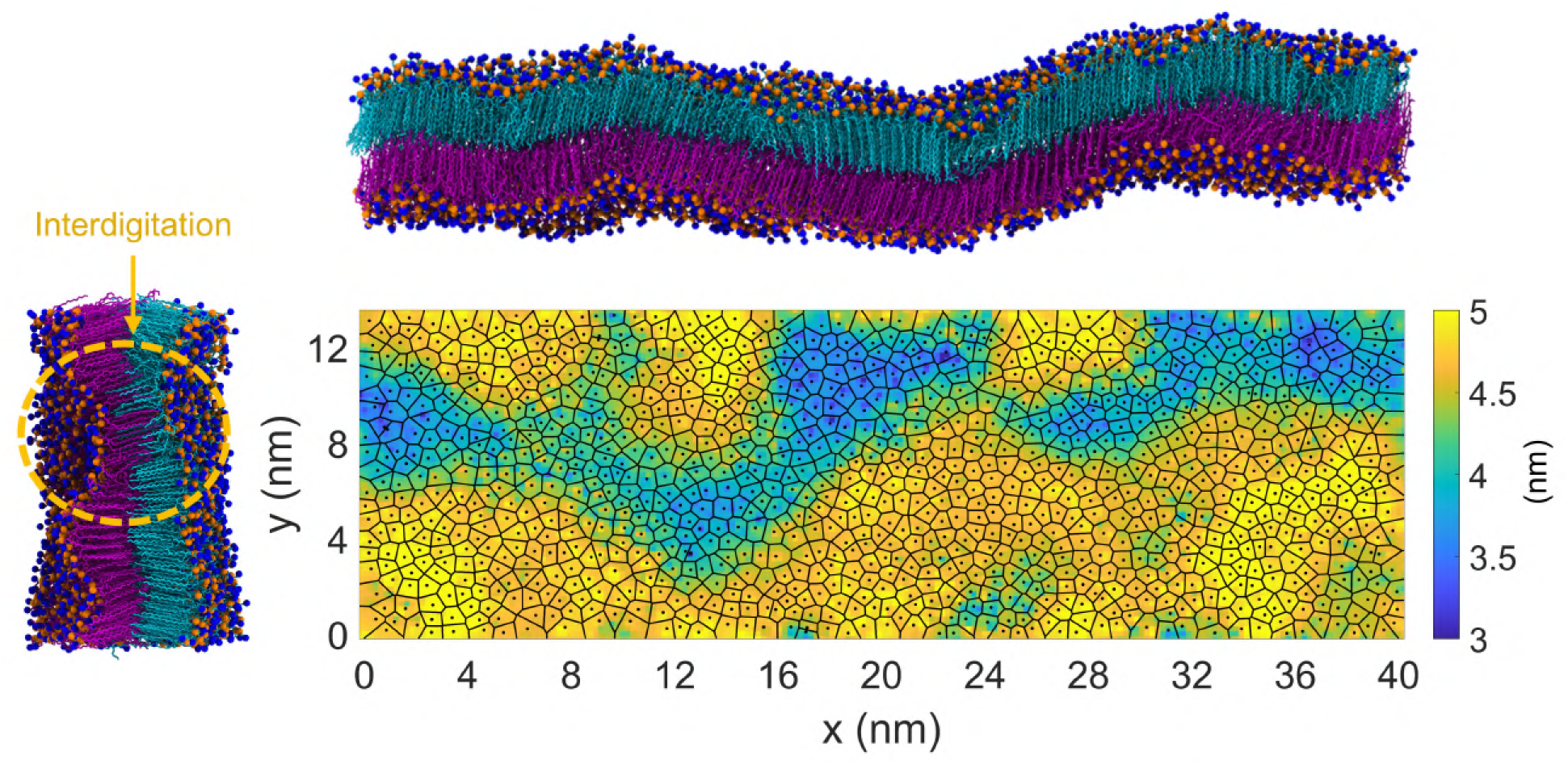
Independent all-atom simulation replicate also shows ripple formation in a large DPPC membrane. Similar to Figure 3, the final snapshot (front view) of a large DPPC membrane after being cooled to 290 K, and its corresponding Voronoi-thickness map, show the formation of symmetric, and asymmetric-interdigitated, ripple structures along the X-axis (shorter box dimension) and Y-axis (shorter box dimension), respectively. Note that the non-interdigitated symmetric ripple along the X-axis is indiscernible from the Voronoi-thickness map.

**Figure S9.**
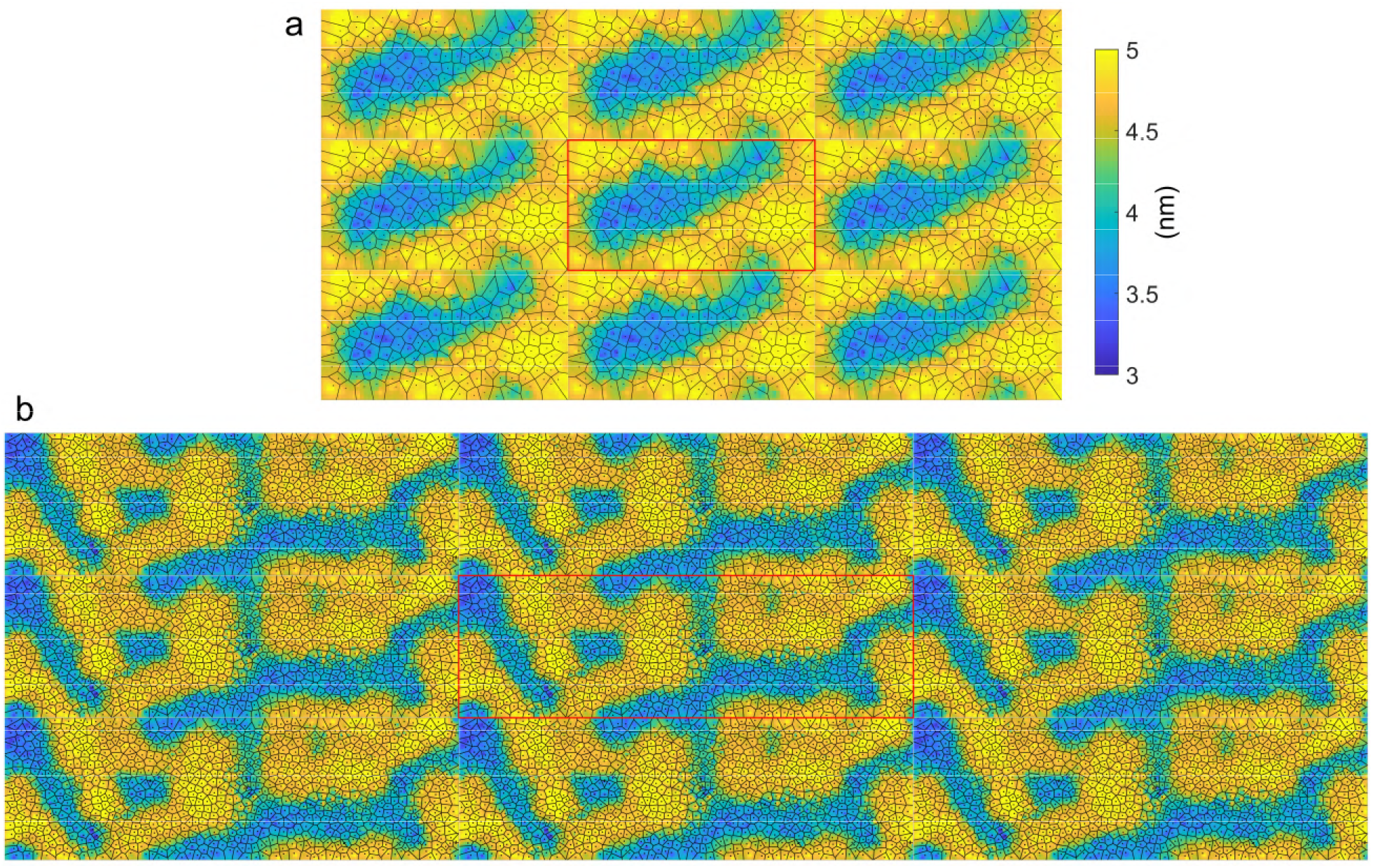
Periodically repeated Voronoi-thickness modulation heatmaps of atomistic DPPC membranes. Voronoi-thickness modulation heatmaps after simulated cooling of atomistic **(a)**368-lipid (Figure 1), and **(b)**2208-lipid (Figure 3) DPPC membranes are shown with periodic images (original simulation box in the middle shown with a red border; same thickness colorbar applies for both systems). The reticulated interdigitated domains in the larger membrane contrast with the island-like isolated domains in the smaller membrane.

**Figure S10.**
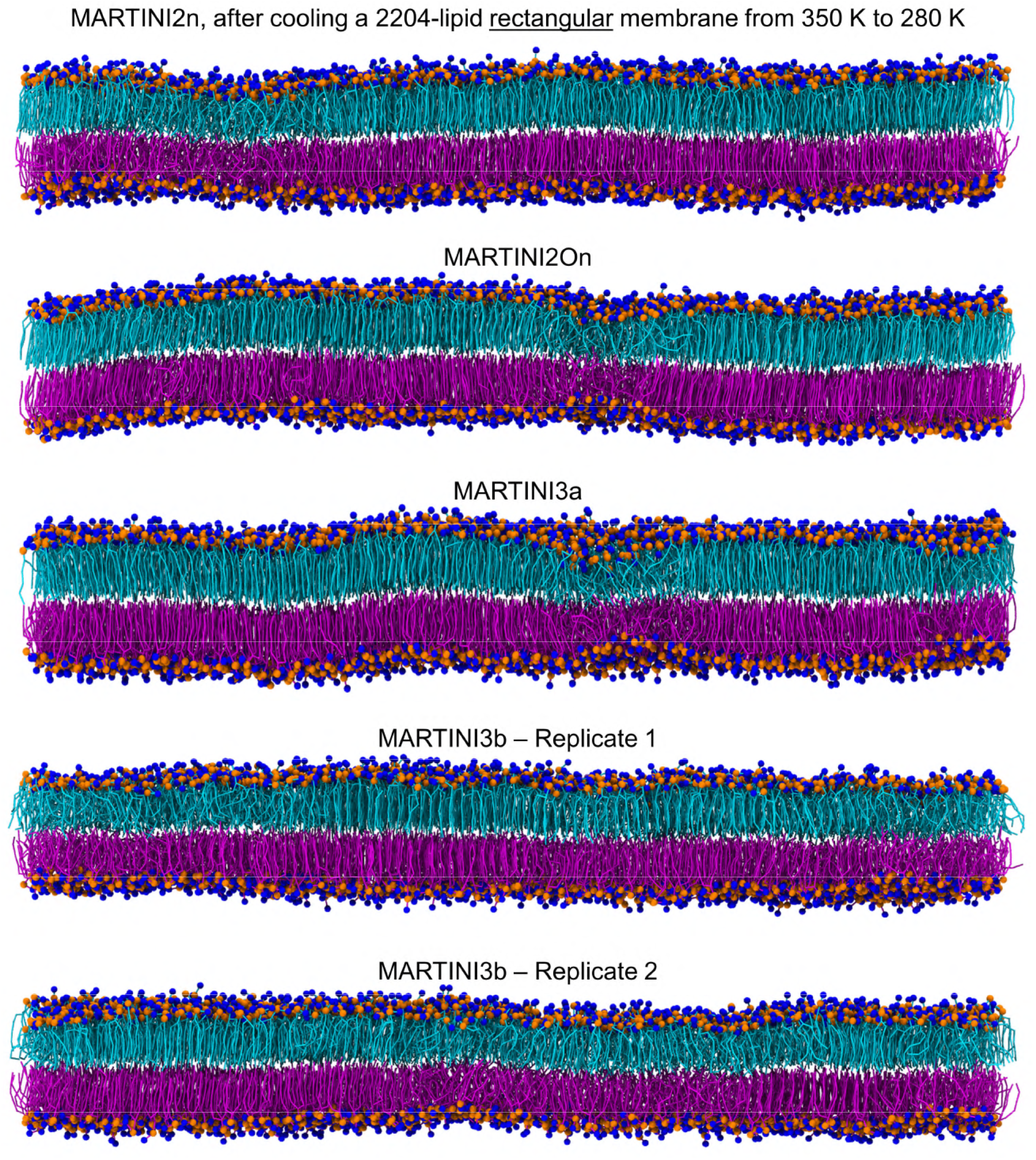
Independent MD replicates confirm the lack of ripple formation in large MARTINI DPPC membranes. Independent MD replicates similar to Figure 4, where large, rectangular, MARTINI2n, MARTINI2On, MARTINI3a, and MARTINI3b DPPC membranes, initially pre-equilibrated in the fluid phase at 350 K, exhibited direct transformation into the straight gel phase upon cooling to 280 K without forming either interdigitated or symmetric ripple-like structures (unlike the behaviour for smaller MARTINI membranes in Figure 2). Note: these membranes were not subject to further EQMD after cooling, and thus a few minor domains remain in the *L*_α_ state. These are expected to disappear upon further equilibration.

**Figure S11.**
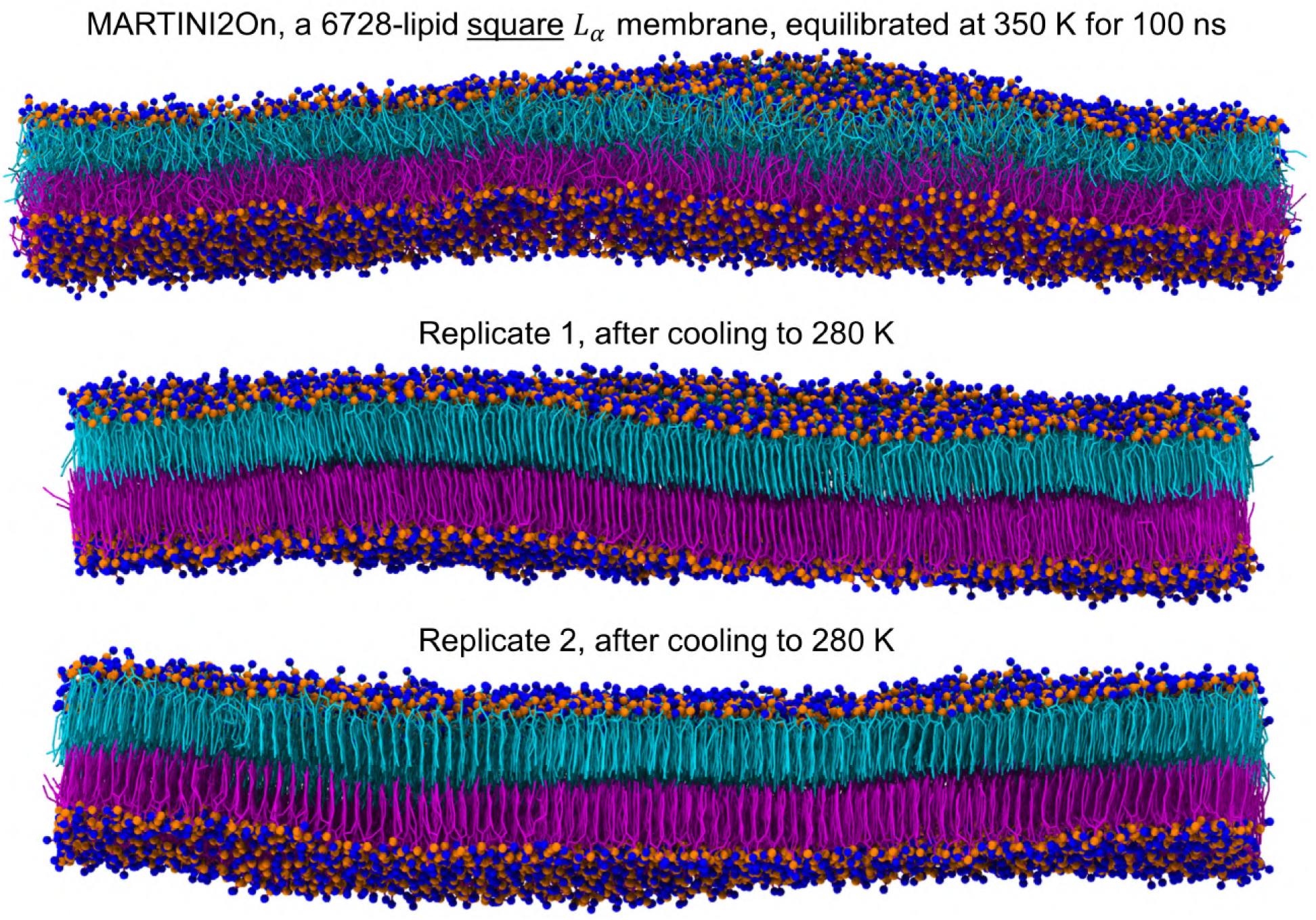
Large square MARTINI DPPC membranes also exhibit a lack of ripple formation. The influence of membrane geometry on ripple formation was also tested by cooling two large, square, MARTINI2On membrane replicates, from 350 K to 280 K. Similar to the rectangular membrane patches in Figure S10, both square membrane replicates directly froze into the gel phase without forming ripple membranes.

**Figure S12.**
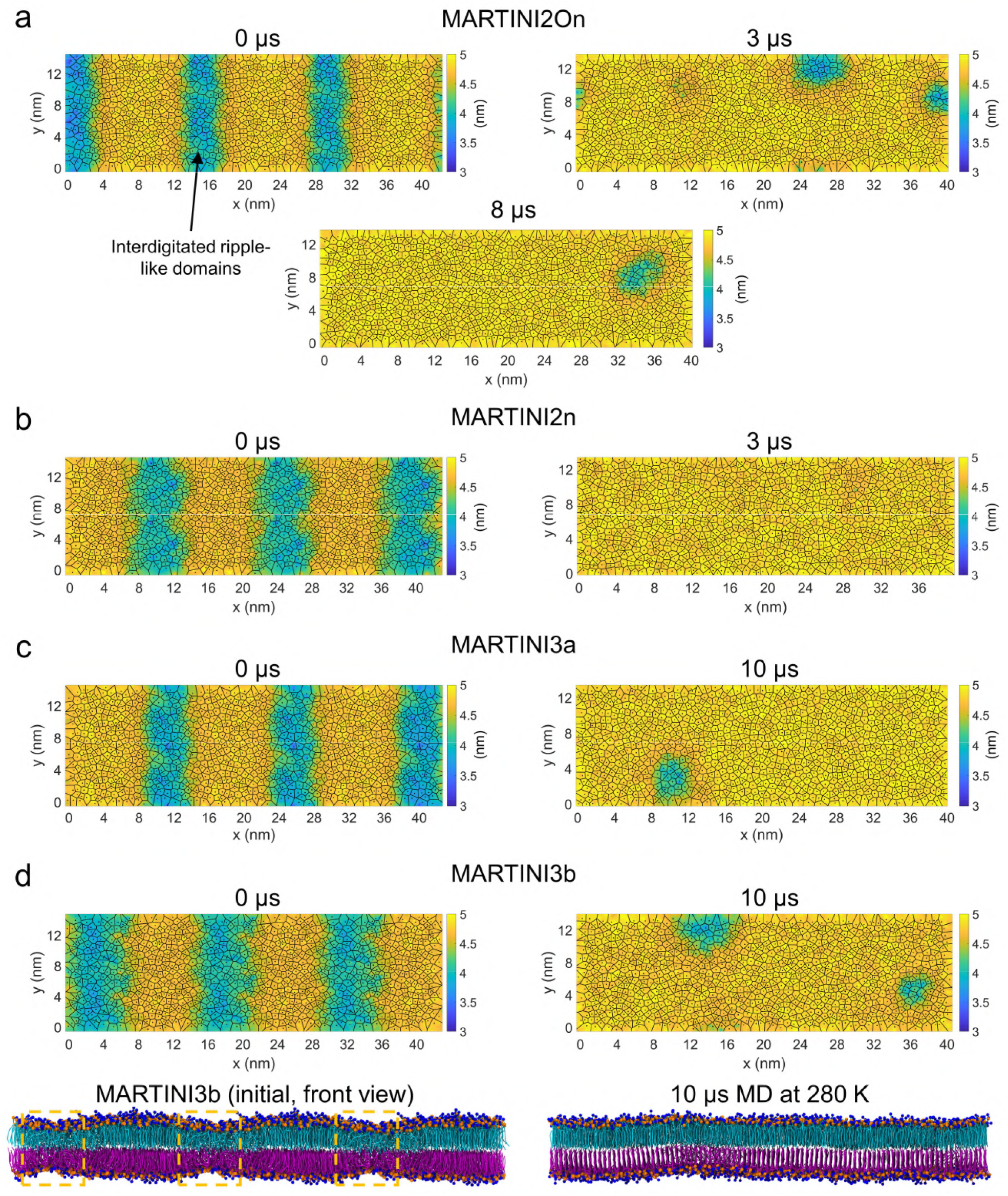
Voronoi-thickness maps from MD of large MARTINI membranes. Voronoi-thickness map from constant temperature MD of large **(a)** MARTINI2On, **(b)** MARTINI2n, **(c)** MARTINI3a, and **(d)** MARTINI3b membranes at 280 K, corresponding to MD snapshots in Figure 5 (snapshots for MARTINI3b shown here). Simulations starting with a periodic ripple-like configuration (0 μs), shows progressive loss of the interdigitated ripple-like domains and formation of the gel phase by 8 μs and 10 μs for the MARTINI2On and MARTINI3a/b membranes, respectively. Notably, the MARTINI2n membrane shows complete loss of interdigitation by 3 μs. While the blue regions at the initial time points (0 μs) correspond to interdigitated domains, they could correspond to either metastable interdigitated (continuously decreasing), or non-interdigitated but partially melted domains, at the final time points. This distinction can be made by simultaneously comparing the MD snapshots (Figure 5 for MARTINI2On, MARTINI2n, MARTINI3a, and above for MARTINI3b) with the Voronoi-thickness maps above.

**Figure S13.**
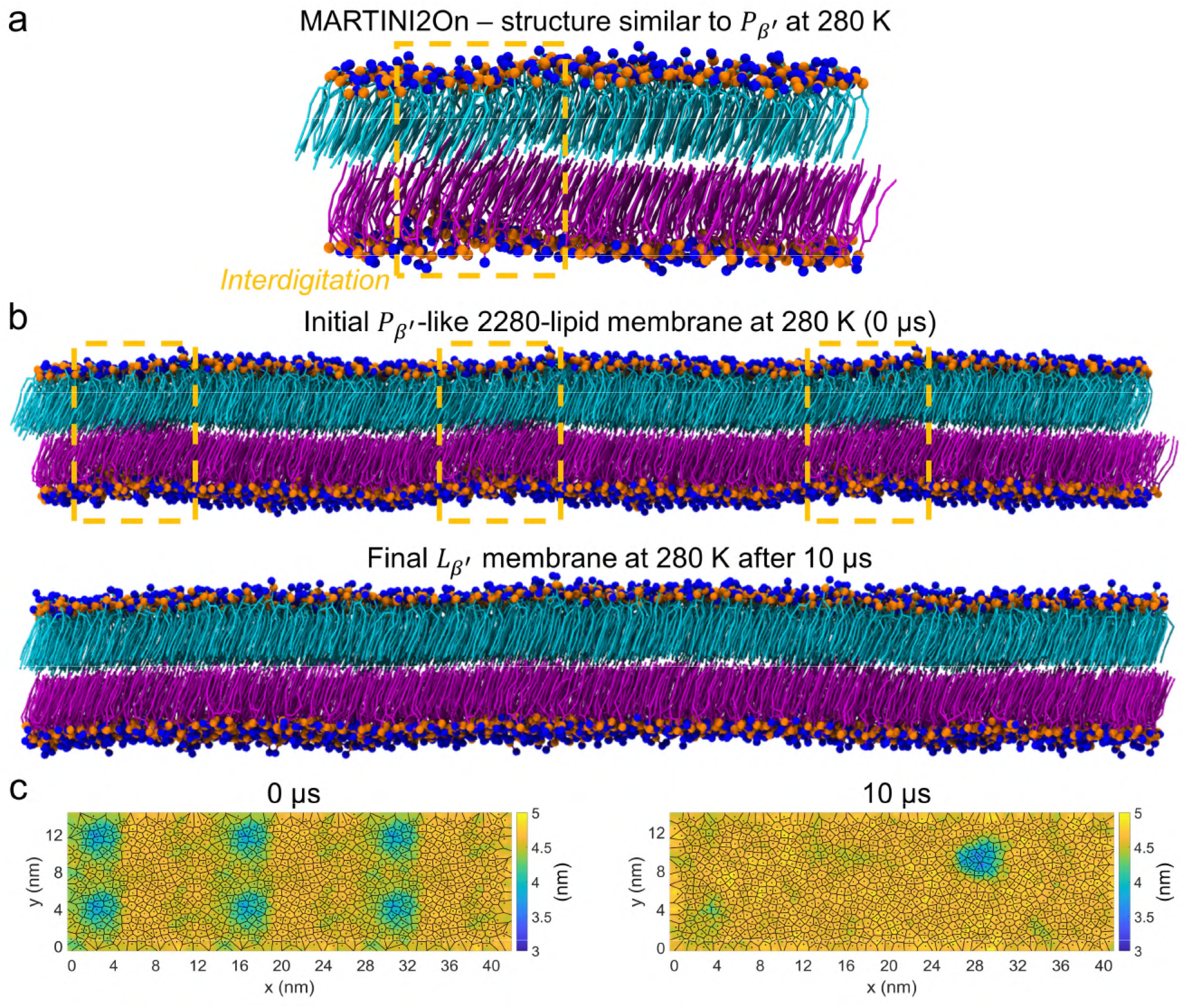
Tilted ripple-like state of MARTINI2On membranes is also a finite size artefact. Simulated cooling of replicate 3 with the MARTINI2On FF (from Figure S3) showed the formation of an interdigitated domain in an otherwise *L*_β′_ membrane at 280 K. This structure was enhanced upon performing constant temperature MD for 5000 ns: final snapshot shown in **(a)**. **(b)** To explore whether this was a finite size effect or a true ripple phase, the smaller membrane was replicated 6x across periodic boundaries to create a larger membrane with repeated ripple-like domains (snapshot at 0 μs), similar to Figure 5*a*. After 10 μs constant temperature MD, the interdigitated ripple-like regions were greatly diminished, thus confirming that the ripple-like state in (a) was indeed a finite size artefact. **(c)** Voronoi-thickness maps pertaining to MD snapshots in (b), where regions of lower thickness in blue indicate the ripple-like domains.

